# Amyloplasts are necessary for full gravitropism in thallus of *Marchantia polymorpha*

**DOI:** 10.1101/2024.11.20.624610

**Authors:** Mimi Hashimoto-Sugimoto, Takuya Norizuki, Shoji Segami, Yusaku Ohta, Noriyuki Suetsugu, Takashi Ueda, Miyo Terao Morita

## Abstract

Gravitropism is a response in which plants sense gravity and determine the direction of organ growth and development. This trait is important for adaptation in land plants. The molecular mechanisms of gravitropism have been studied mainly in flowering plants, but there is limited research on other organisms. In this study, we examined the gravitropic response of the liverwort *Marchantia polymorpha*, a model for investigating the evolution of land plants. We found the tips of the thallus extend upward and form several straight narrow structures in the dark. These growth directions were always in the opposite direction of gravity, and clinostat treatment disrupted them. The parenchymatous cells in the narrow structures contained amyloplasts, and the sedimentation of the amyloplasts preceded the gravitropic curvature, suggesting their role as statoliths. The starchless mutants, Mp*pgm1 and* Mp*aps1* were generated, and an abnormal direction of growth was observed in the narrow structures, but they tended to elongate upward. These observations indicate that although amyloplasts are required for complete gravitropism, plants can sense gravity without well-developed amyloplasts. These results suggest that land plants use amyloplasts as statoliths but also have amyloplast-independent mechanisms of gravitropism. These results suggest that land plants use amyloplasts as statoliths but also have amyloplast-independent mechanisms of gravitropism.

**Highlight:** In *M. polymorpha,* amyloplasts act as statoliths in parenchyma cells and are important in gravitropism but it was not completely lost without starch granules.

## Introduction

Plants are essentially sessile organisms and can sense various environmental stimuli and then adjust their growth direction to a position more suitable for water absorption, photosynthesis, and reproduction. Gravity and light are the most important directional environmental stimuli controlling growth direction (Knight, 1806; Hangarter, 1997; Kiss, 2000). Different from other environmental stimuli, gravity is continuously presented, so plants can use this reliable indicator of their direction even in dark conditions. Gravitropism in plants is the development or growth of plant organs in a specific direction in response to gravity, which is composed of four sequential steps: gravity perception, signal formation, signal transduction, and differential growth of the upper and lower tissues of the responding organ (Tasaka et al., 1999; Morita and Tasaka, 2004; Morita, 2010).

In various stem and root tissues of flowering plants, the gravity susceptors are believed to be sediment in specialized cells, which are thought to be involved in gravity sensing (Haberlandt, 1900; Némec, 1900). A pile of studies has supported the concept of starch statoliths theory in the gravity perception in gravitropism (Sack, 1991; Konings H, 1995; Blancaflor and Masson, 2003).

Physiological and genetic investigations have provided evidence that amyloplasts, starch-filled plastids, are necessary for full gravitropic sensitivity of the root and the shoot in *A. thaliana*. *pgm* (*plastidial phosphoglucomutase*) mutant with impaired plastidic phosphoglucomutase activity is a starch-deficient mutant of *A. thaliana*, and shows reduced gravitropism (Caspar and Pickard, 1989; Kiss et al., 1989; Kiss et al., 1997). Other starchless or starch-reduced mutants that contain immature amyloplasts show reduced gravitropism in both the shoot and the root (Kiss *et al*., 1989, 1996, 1997; Weise and Kiss, 1999). The extent of gravitropism is positively correlated with the starch content of the amyloplast, suggesting that the mass of the amyloplast indeed affects the magnitude of the gravitropic response (Kiss *et al*., 1996).

The statoliths in the alga *Chara* sp. living in water, are vesicles filled with aggregates of barium sulfate crystals (Sievers *et al*., 1991; Braun and Sievers, 1993; Hodick *et al*., 1998). Much of the early work on the effects of gravity on bryophytes was summarized by Petschow (Petschow, 1933). He found that some moss and liverworts had intracellularly sedimented starch granules in their thalli and grew upward in the dark. Furthermore, amyloplast sedimentation and upward curvature also appeared in single-cell protonemata of mosses, suggesting a correlation between the perception of gravity and settled statoliths (Petschow, 1933; Sack *et al*., 2001). However recent work revealed that rhizoids of a model moss *Physcomitrella patens* (*Physcomitrium patens*) exhibit gravitropism but did not find any detectable starch granules (Zhang et al. 2019). It remains unclear whether the starch granules are required as gravitropic signaling factors in bryophyte, partly because molecular genetic analysis was poorly developed. *M. polymorpha,* a liverwort, has recently been established as a model plant species with available genomic information (Bowman *et al*., 2017; Bowman, 2022) and various tools for molecular genetic analysis such as CRISPR/Cas9-based genome editing (Sugano et al., 2018; Kohchi et al., 2021). In this study, we focused on the gravity responsiveness of thalli of *M. polymorpha* and investigated its basic characteristics. In addition, genetic and physiological analyses were conducted to understand the role of starch grains in the response.

## Materials and methods

### Plant Material and Growth Conditions

Takaragaike-1 (Tak-1) of *Marchantia polymorpha* was used as a wild type and genetic background for transgenic lines. To observe MpPGM1-Citrine, transgenic plants expressing *_pro_*Mp*EF1α:*Mp*PGM1-Citrine* (Norizuki *et al*., 2023) were used. Gemmae of *M. polymorpha* plants were horizontally grown on half-strength Gamborg B5 medium supplemented with 1% (w/v) sucrose (pH 5.5, 1% agar) (Gamborg *et al*., 1968) at 22°C under continuous light. To generate narrow structures from the original thallus, plates of the light-grown thalli were placed vertically in darkness. After some elongated portions of the thallus emerged, the plates with plants on them were turned 90° or 180°. The experiment to determine if a part of the thallus buried in the soil would come out of the ground was conducted as follows. 2 week-old light-grown thalli were covered with the soil of 1: 1 (v/v) mixture of vermiculite and Metromix 350 (Scotts-Sierra Horticultural Products, U.S.A.) in 1 cm deep and then grown with 5000-fold diluted Hyponex solution (Hyponex, Japan) applied as nutrient for 5 weeks at 22°C under continuous light.

### Identification of MpPGMs, MpAPSs, and MpAPLs

Amino acid sequences of Mp*PGM*1 (Mp4g13750.1/Mapoly0202s0014.1), Mp*PGM2* (Mp5g10560.1/ Mapoly0048s0016.1), Mp*APS1* (Mp1g15530.1/ Mapoly0033s0108.1), Mp*APL1* (Mp2g11530.1/Mapoly0023s0119.1), Mp*APL2* (Mp4g20740.1/Mapoly0101s0020.1), and Mp*APL3* (Mp4g10120.1/Mapoly0132s0055.1) were obtained from *M. polymorpha* genome version 5.1 (Montgomery *et al*., 2020) in MarpolBase (http://marchantia.info/) using these proteins of *A. thaliana* as queries. A domain search was performed using SMART (http://smart.embl-heidelberg.de/) (Letunic and Bork, 2018; Letunic *et al*., 2021). Transit peptides of MpPGM1 and MpAPS1 were deduced using ChloroP 1.1 Server (http://www.cbs.dtu.dk/services/ChloroP/) (Emanuelsson, et al., 1999). We followed the nomenclature by Bowman et al. (2016) for nomenclature of genes, proteins, and mutants of *M. polymorpha* (Bowman *et al*., 2016).

### Phylogenetic analysis of PGMs, APSs/APLs

Phylogenetic analyses of PGMs and APSs/APLs were according to Norizuki et al. (Norizuki *et al*., 2019). As a substitution model, we used the WAG+G+I (for PGM) or the LG+G+I+F (for APS APL) model which was selected by Smart Model Selection in PhyML (Lefort *et al*., 2017). Bootstrap analysis was performed by resampling 1,000 sets. The sequences used in the phylogenetic analysis were included in Supplementary Material S1.

### Construction and transformation

To construct the *pro*Mp*EF1*α:Mp*APS1-Citrine*, open reading frames of Mp*APS1* were amplified by polymerase chain reaction (PCR) from cDNA obtained from thalli of *M. polymorpha* accession Tak-1 using the oligonucleotides MpApS1_Fw and MpApS1_Rv, and the amplified product was subcloned into pENTR^TM^/D-TOPO (ThermoFisher, USA) according to the manufacturer’s instructions. The resultant sequence was then introduced into pMpGWB308 (Ishizaki *et al*., 2015) using the Gateway LR Clonase^TM^ II Enzyme Mix (ThermoFisher) according to the manufacturer’s instructions. To construct CRISPR/Cas9 vectors, two complementary oligonucleotides in the sequences of Mp*PGM1* (MpPGM1_gDNA_Fw and MpPGM1_gDNA_Rv), and Mp*APS1* (MpAPS1_gDNA_Fw and MpAPS1_gDNA_Rv) were synthesized and annealed, and the resulting double-stranded fragments were subcloned at the *Bsa*I site of the pMpGE_En03 vector (Sugano *et al*., 2018) using the DNA ligation kit Ver.2.1 (Takara Bio, Japan) according to the manufacturer’s instructions. The resultant guide RNA cassette flanked by the *att*L1 and *att*L2 sequences in pMpGE_En03 was then introduced into the pMpGE010 vector (Sugano *et al*., 2018) using the Gateway LR Clonase^TM^ II Enzyme Mix. The primer sequences used in this study are listed in Supplementary Table S1. The transformation was performed as previously described (Kubota *et al*., 2013). Transformants were selected on plates containing 10 mg/L hygromycin B and 250 mg/L cefotaxime for the pMpGE010 vectors and 0.5 μM chlorsulfuron and 250 mg/L cefotaxime for the pMpGWB308 vector.

### Genotyping

For the genotyping of mutants generated by CRISPR/Cas9, genome DNA was extracted from thalli by the extraction buffer (1 M KCl, 100 mM Tris-HCl (pH 9.5), and 10 mM EDTA). Genome regions of Mp*PGM1* and Mp*APS1* were amplified by PCR using KOD FX Neo (TOYOBO, Japan) using the oligonucleotides MpAPS1_GT1_Fw, MpApS1_Rv, MpPGM1_GT1_Fw and MpPGM1_GT2_Rv, and mutations in these genome fragments were analyzed by direct sequencing using the oligonucleotides MpAPS1_seq1_Fw and MpPGM1_seq1_Fw.

### Clinostat rotation

Thalli were grown on a plate in light for 2 weeks. The plate was covered with an aluminum foil to culture thalli in darkness and equipped with a 3D-Clinostat for 2 weeks (Plant Gravity Response Research Equipment; Kitagawa, Japan).

### Time-lapse images

As described above, 2-week light-grown thalli were placed on vertically-held plates in the dark for 2 weeks to form the narrow structures. The plates were then turned 90° to a horizontal position. These developments of narrow structures were recorded continuously in darkness with a time-lapsed video image recording camera (Brinno TC200 Pro; Brinno, Taiwan) with a low destruction lens (VS-LD4; VS Technology, Japan), using infrared radiation with a peak emission wavelength of 950 nm as the monitoring light (KB850-7224LE KS-602983; KOSU system, Japan).

### Lugol’s Iodine Staining

Thalli with protruding narrow structures were fixed in 80% (v/v) ethanol for 30 min under a vacuum at room temperature. The fixed tissues were washed twice for water and stained in 0.05 mol/L iodide solution (Wako). This method did not remove the weak dark pigment (Supplementary Fig. S1A-C), so ClearSee solution was used before iodine treatment to more clearly compare staining differences between starch-deficient mutants and the wild-type (Kurihara et al., 2015; Segami et al., 2018). Samples were fixed in 4% (w/v) paraformaldehyde in phosphate-buffered saline (PBS) for 30 min under a vacuum at room temperature. The fixed tissues were washed twice for 1 min in PBS and cleared with ClearSee version1 (10% (w/v) xylitol, 15% (w/v) sodium deoxycholate, and 25% (w/v) urea). Sodium deoxycholate has a weak decolorizing activity for iodine staining, so the samples were transferred to 10% (w/v) xylitol and 25% (w/v) urea. Finally, the tissues were stained in 0.05 mol/L iodide solution. Sample images were obtained by a digital microscope Olympus DSX510 (Olympus). modified pseudo-Schiff-propidium iodide (mPS-PI) Staining

Narrow structures of thalli were fixed in fixative (50% (v/v) methanol and 10% (v/v) acetic acid) for at least 16 h. The fixed tissue was rinsed with water and incubated in 1% (v/v) periodic acid at room temperature for 40 min. The tissue was rinsed again with water and incubated in Schiff reagent with propidium iodide (100 mM sodium metabisulphite and 0.15 N HCl; propidium iodide to a final concentration of 100 mg/mL was freshly added) for 3 h in darkness. The samples were washed twice in water and then transferred onto microscope slides. Several drops of Hoyer’s solution (15 g gum arabic, 100 g chloral hydrate, 10 g glycerol, and 25 mL water) were placed over the tissue, and a cover slip was placed on top.

### Microscopy

Bright-field images of Lugol-stained samples were obtained using a digital microscope (DSX110; Olympus). For confocal microscopic observation of thallus cells expressing MpPGM1-Citrine or MpAPS1-Citrine, five-day-old thalli were observed with an LSM780 confocal microscope (Carl Zeiss) equipped with an oil immersion lens (×63, numerical aperture = 1.4) as described previously (Norizuki et al. 2023). CLSM observations for mPS-PI-stained samples were conducted with an upright FV1000 CLSM (Olympus). The excitation wavelength was 559 nm and the transmission range for emission was 655-755 nm. The images were obtained using Z-stack (consecutive 5 µm thick optical slices) with Olympus FluoView software. Optimal depth was calculated by the numbers of the stacks.

### Data analysis

Direction and length of narrow structures, manual tracking of the tip of narrow structures, and distribution of amyloplasts in a cell were measured and analyzed with open-source software Fiji (ImageJ 1.53q, Java 1.8.0) (Schindelin *et al*., 2009).

## Results

### Protruded narrow structures emerged from dark-grown thallus in M. polymorpha and grew upward

To eliminate the influence of light and to examine the gravity response in thalli of *M. polymorpha*, thalli were grown in the dark. Gemmae of the wild-type plant (Tak-1) were placed on agar plates containing sucrose and grown horizontally under light for 2 weeks (Fig. 1A). Then they were placed vertically in the dark for 2 weeks to examine for morphological changes in response to gravity. We found that thin structures (hereafter we call them narrow structures) came out from the thalli and grew straightly upward (Fig. 1B). Then we tracked their growth with time-lapse imaging (Supplementary

**Fig. 1.**
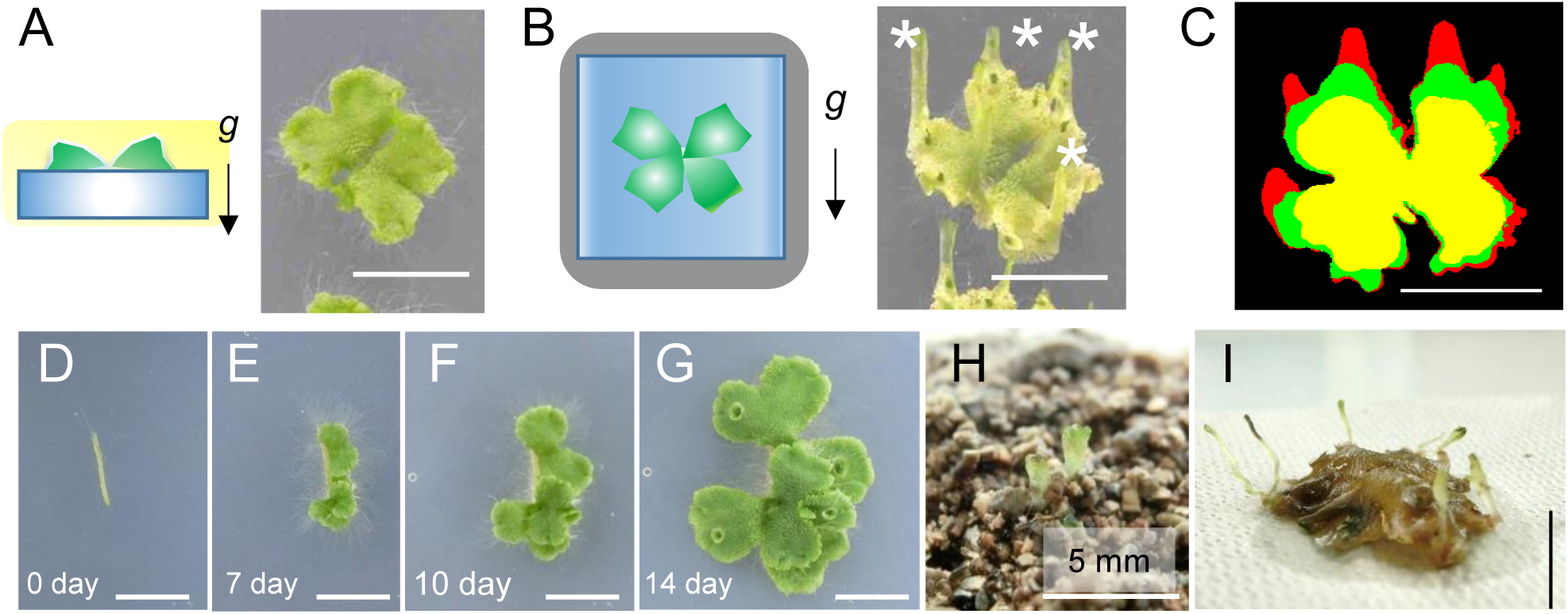
*M. polymorpha* formed narrow structures growing upright in the dark. 2-week-old wild-type (Tak1) thalli grown in the light (A) were placed for an additional 2 weeks in a vertical position in the dark (B) shown with diagrammatic representations. The black arrows indicate the direction of gravity. (C) Representative morphological changes of thalli transferred to the dark. The shape of the thallus at 11 days of light grown was shown in yellow, 3 and 7 days after transition to the dark were shown in green and red, respectively. (D-G) Dissected narrow structure of a thallus grown in the dark for 2 months was transferred to the light. This produced some radial tissue and formed a thalli-like shape in the light. The number of days after light exposure is shown in the lower left. (H, I) 2-week-old light-grown thalli were buried in the soil at a depth of 1 cm. Narrow structures appeared on the surface of the soil (H) and excavated whole plant (I) 5 weeks after it was buried in the ground. Scale bars = 1cm in A-G, and I; 5 mm in H.

Movie S1). When 11-day-old thalli grown under light were placed in the dark (Fig. 1C, 0 day; Yellow), the tips of the upper parts swelled to form bulges on the third day (Fig. 1C, 3^rd^ day; Green). Then, the elongation of the narrow structures accelerated further, and the narrow structures continued to grow upward (Fig. 1C, 7^th^ day; Red). In the time-lapse movie, we found that the upper part of the narrow structures appeared to grow faster than the lower part of each thallus (Supplementary Movie S1). Presumably, it is not as easy as the upper part growing upwards, as the lower part must bend its body to grow upwards (Supplementary Movie S1).

A bulge seemed to form from an apical notch of the thallus at the beginning of the formation process of the narrow structure (Supplementary Fig. S2A). It was elongated in the dark, and the tip of the narrow structure had two apical notches (Supplementary Fig. S2B). Gemma cups were produced on the narrow structures and produced gemma normally, although the cups were very shallow (Supplementary Fig. S2C, D). The rhizoids on the ventral side and the gemma cups on the narrow thallus’s dorsal side indicate a clear dorsiventral pattern (Supplementary Fig. S2D). Furthermore, when the narrow structure was grown again under light irradiation, some parts grew into a shape resembling a thallus (Fig. 1D-G). These results indicate that the narrow structure possesses features of the original thalli.

If the plants were buried in the soil, would they sprout? We buried 2-week-old thalli 1 cm deep in the soil to shade out the light, and grew in the soil (Supplementary Fig. S3). After 5 weeks, we found that some small parts of thalli emerged from the soil (Fig. 1H). When we dug it out, we found that the size of the original thalli had not changed much (Supplementary Fig. S3C, D), but they had lost their green color, whereas newly emerged narrow structures remained green (Fig. 1I, Supplementary Fig. S3D). It was similar to what was observed in the plate experiments in the dark (Fig. 1B). The extension of part of the thallus above the ground in the dark conditions may be an adaptive trait that allows *M. polymorpha* to survive when buried by obstacles.

### Narrow structures can sense changes in the direction of gravity

We investigated whether the direction of elongation of the narrow structures depends on the direction of gravity. The 7-day-old plants were dark-treated for 12 days to induce the formation of narrow structures (Fig. 2A). When the plants were rotated 90° for 4 days, the narrow structures grew in the direction opposite to that of gravity (Fig. 2B). After another 90° rotation for 4 days, the narrow structures elongated in the direction opposite to gravity again (Fig. 2C).

**Fig. 2.**
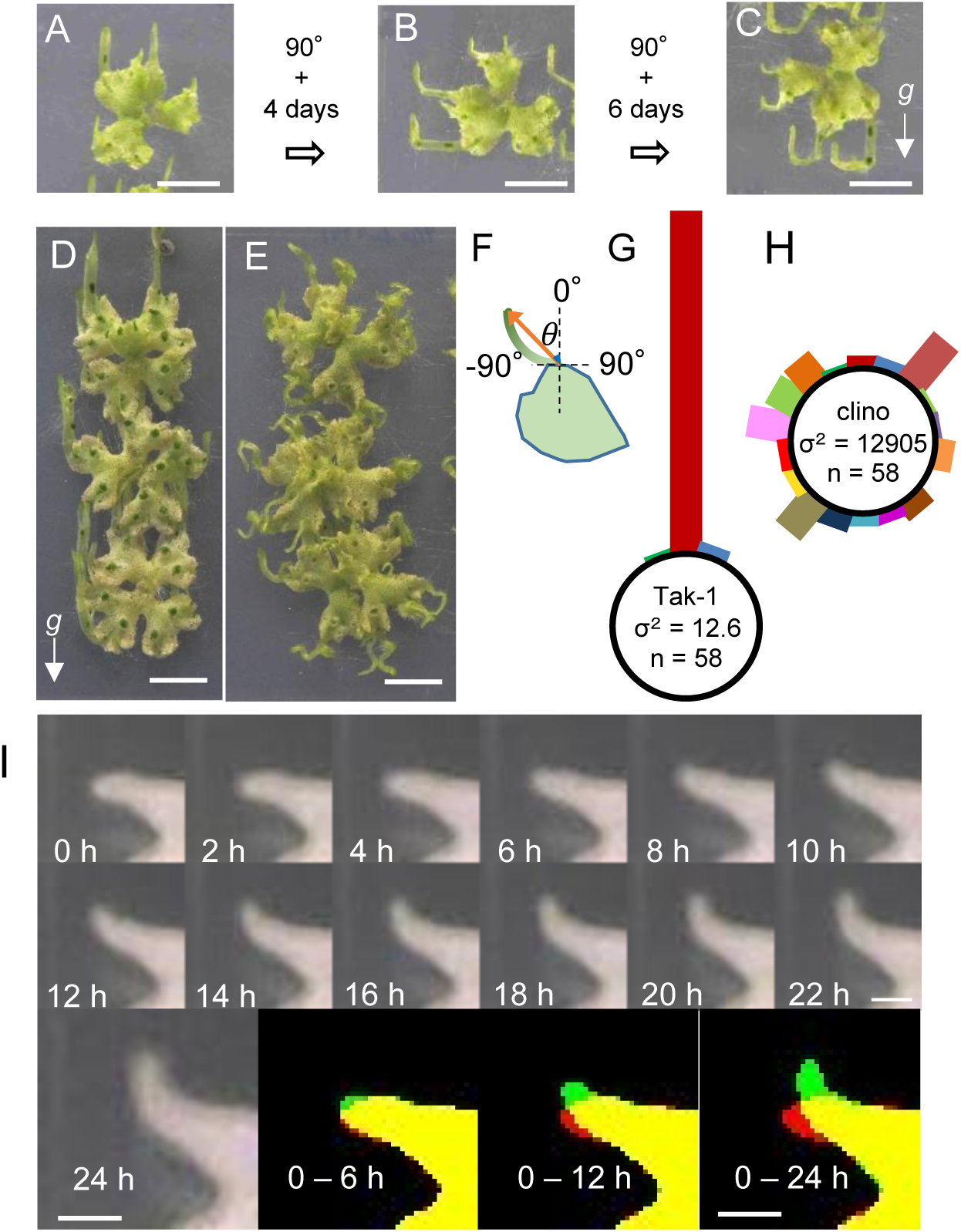
Narrow structures showed a sensitive response to gravity. (A-C) The narrow structures showed negative gravitropism. 7 day-old gemmalings grown in the light were placed vertically in the dark for 12 days (A). The thalli with vertically oriented narrow structures were turned to 90° and grown for 4 days resulting in upright elongation (B). Additional 90° turning and 6 days of growth induced further vertical growth of the narrow structures (C). (D, E) Thalli grown in the light for 2 weeks were grown vertically (D) or on a 3D clinostat (E) in the dark for 2 weeks. (F-H) Angles (0) from the basal position of the narrow structures to the top (G) were measured in vertically grown (G) and clinostat-treated (H) Tak-1. The bar length is proportional to the number of plants observed in each 20° bin. (I) Enlarged images of a tip of the narrow thallus and superimposed images of before (Time 0, red) and 6, 12, and 24 h after 90° rotation (green) are shown. Plants were grown for 10 days in the light and 11 days in the dark vertically and then rotated at 90°. The elapsed time (hour) after the 90° rotation is shown at the top of the images. The superimposed image of before (Time 0, red) and 24 h after 90° rotation (Time 24 h, green) indicates the movement of the narrow thallus. Scale bars = 1 cm in A-E; 2 mm in I.

To further confirm the gravity dependence of the growth direction of narrow structures, we examined that direction when thalli were placed on a 3D clinostat, an experimental device that simulates microgravity environments (Hoson *et al*., 1992). When the thalli were placed vertically in the dark for 2 weeks, they elongated straightly in the direction opposite to that of gravity (Fig. 2D), but when thalli were placed in the 3D clinostat in the dark for 2 weeks, they did not elongate straightly but turned and twisted in various directions (Fig. 2E). We quantified growth angles of narrow structures under gravity on earth (1×g) or on the 3D clinostat by measuring the angle formed between the growing direction of the tip and the vertical axis (Fig. 2D-H). We found that most of the narrow structures of the plants placed erectly grew in the direction opposite to gravity (Fig. 2D, G), on the contrary, thalli on 3D clinostat grew irrespective of the directions of gravity (Fig. 2E, H). There was no statistical difference in the growth (length of emerged narrow structures) between treatments (Supplementary Table S2). These results indicate that narrow structures are sensitive to gravity.

To observe morphological changes in the gravity response of narrow structures in detail, the plant with narrow structures grown in the dark for 11 days was rotated 90°, and time-lapse images were taken every 2 hours (Fig. 2I, Supplementary Movie S2). All of the narrow structures bent upward after 24 h (Fig. 2I, 0-24 h, Supplementary Movie S2). Detailed observation showed that the tip of the narrow structure changed its direction of growth approximately 6 hours after gravistimulation and grew in the direction opposite to gravity after 12 hours, and the tip turned straight up after 24 hours under our experimental conditions (Fig. 2I; 0-6 h, 0-12h and 0-24 h). This result indicates that response to gravity began to be observed 6 hours after gravistimulation, and the direction of elongation was determined until 24 h. After 24 h of gravistimulation, the position of the tip was shifted from the original elongated thallus, suggesting that differential growth of the narrow structure, as well as cell proliferation, causes morphological changes. Therefore we concluded the gravity response in the elongated thallus of *M. polymorpha* is negative gravitropism.

### Amyloplasts sediment in cells of narrow structures

To investigate the involvement of starch granules in the gravitropism of the narrow structure, we performed Lugol’s staining with potassium iodide. Seven-day-old thalli were placed in the dark for 2 weeks, and the whole plants were treated with Lugol’s solution. The results showed that only the narrow structures protruding from the original thallus were stained (Supplementary Fig. S1A). This indicates that starch granules were present only in the narrow structures but not in the original thallus. The narrow structures extended from the original thalli showed stronger staining at the tip (Supplementary Fig. S1B). Staining was stronger closer to the tip, with staining becoming sparser and lighter away from the tip. For each cell, more intense staining was observed on the gravity side, implying the stained structures are starch-containing amyloplasts (Supplementary Fig. S1C).

The modified pseudo-Schiff-propidium iodide staining (mPS-PI) (Haseloff, 2003; Flütsch et al., 2018; Zhang et al., 2019)was performed to observe the starch-containing amyloplasts in each cell in detail (Fig. 3). On the dorsal epidermis, small intracellular granules were uniformly detected, and these seemed undeveloped amyloplasts (Fig. 3A, Surface). There was parenchymatous storage tissue under one layer of epidermis with air chambers (Shimamura, 2016). We found large amyloplasts settling at the bottom of the cells in the parenchymatous storage tissues. The cells in the parenchymatous storage tissue were not simply layered, but rather slightly intricate, and became larger as the depth increased (Supplementary Fig. S1D-F). We decided to examine the cells at a depth of 40 µm inside from the surface calculated by z-stack images, where the cells were large enough and the sedimentation of amyloplasts was relatively easy to observe (Fig. 3A, 40 μm inside, Fig. S1F). At the very close to the tip of a narrow structure, the amyloplasts were packed tightly in the small cells (Fig. 3A, 200-400 µm), however, at about 500-700 µm from the tip, the cells became larger at the same depth, and clear sedimentation of amyloplasts with strongest signals was observed (Fig. 3A, 500-700 µm). At further distances from the tip, the cells became larger and many of the amyloplasts settled on the gravitational side of the cell (Fig. 3A, 800-1000 µm). Quantification of the location of the amyloplast is shown in Fig. 3B. The cells were divided into four sections (I-IV) in the direction of gravity, and the total areas of amyloplast exclusivity within each section were examined (Fig. 3B, left) and we calculated “amyloplast occupancy” as the rates of the area occupied by amyloplasts in each section (Fig. 3B, right). Small granules were uniformly detected in all areas of the epidermal cells regardless of the distance from the tip of the narrow structures (Fig. 3B, Surface). On the contrary, greatest number of amyloplasts was detected in region IV at a depth of 40 µm in the range 200-1000 µm from the tip compared to other areas, indicating that amyloplasts settle in the direction of gravity (Fig. 3B, 200-400 µm, 500-700 µm, 800-1000 µm; Kruskal-Wallis rank sum test followed by Steel-Dwass, P<0.001, N > 8). These findings indicate that the amyloplasts in *M. polymorpha* are settling in the direction of gravity specifically in cells of parenchymatous storage tissue.

**Fig. 3.**
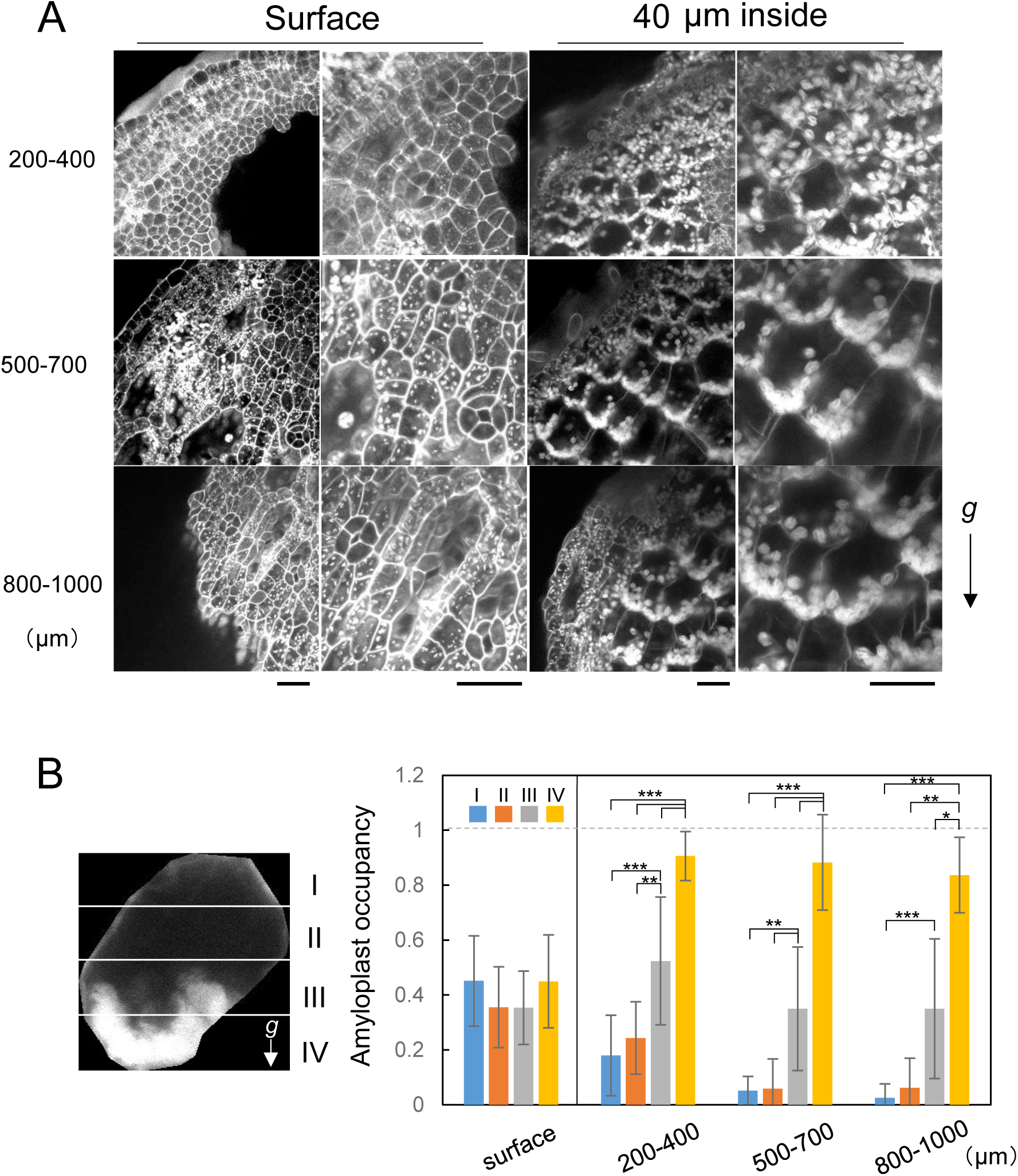
Visualization of starch granules of narrow structures. (A) Starch granule accumulation was evaluated by staining with modified propidium iodide staining. Images on the surface (Left) and 40 μm inside from the surface (Right) of the narrow structures with enlarged images were shown. The numbers on the left of the images indicate the distance from the tip. Scale bars = 50 μm. (B) Statistical analysis of amyloplast distribution. A cell was separated into four regions (I–IV) along the gravity vector (left), and the area of amyloplasts in each region was counted and shown as amyloplast occupancy (%) (right). Each bar indicates the average ± SD (N=8). Significant differences were identified by the Kruskal-Wallis rank sum test followed by Steel-Dwass, and are indicated with * P<0.05, **P<0.01, ***P<0.001.

### Gravistimulation induces amyloplast sedimentation in cells located at the tips of narrow structures before gravitropic curvature

Next, we examined the kinetics of amyloplast relocation in cells of narrow structures upon gravistimulation by 90° reorientation. The amyloplasts began to settle after 3 h, although some were still settling in the original direction of gravity in cells within 700 μm from the tip (Fig. 4A, upper). Six hours later, the amyloplasts settled in the new direction of gravity (Fig. 4A, middle). After 24 hours of gravistimulation, we could find clear amyloplast settlement in most cells near the tip (Fig. 4A, bottom). The relative position of the amyloplasts in a cell was quantified separately for the original (Fig. 4B, 1^st^) and new gravity directions (Fig. 4B, 2^nd^). The results show that after 3 h of 90° gravistimulation, the amyloplast occupancy increased as the area number increased from region I to region IV indicating that new gravity-directed amyloplast sedimentation was taking place. At the same time, that also increased from region V to region VIII, indicating that the original amyloplast sedimentation has not yet been eliminated (Fig. 4C). After 6 h and 24 h, amyloplast occupancy also increased from region I to region IV, but no difference was found in that from region V to region VIII, indicating that there was no bias of amyloplasts to the original direction of gravity (Fig. 4D, E). These results suggest that amyloplasts may act as statoliths in *M. polymorpha*.

**Fig. 4.**
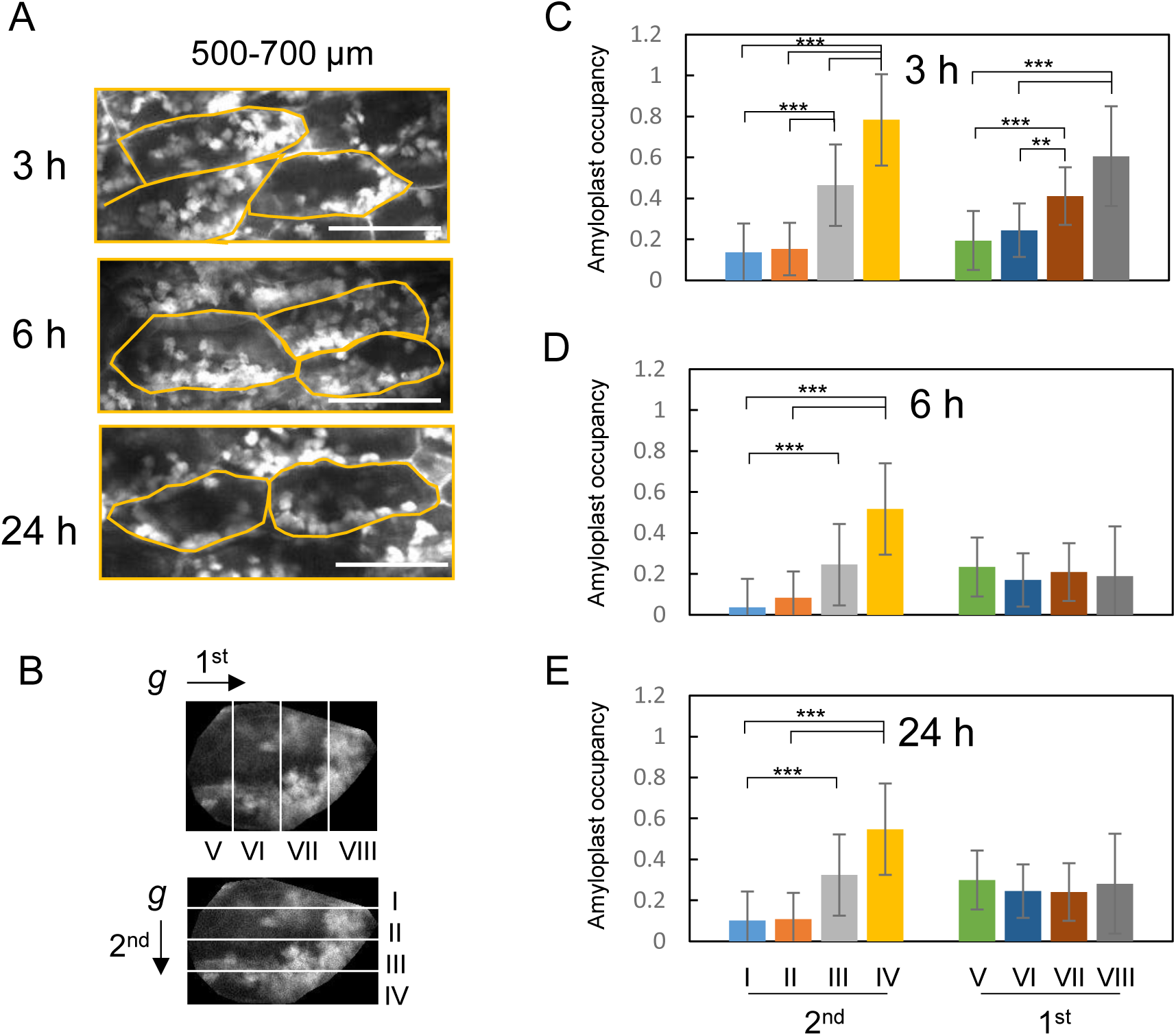
Amyloplast sedimentation and growth direction change of narrow structures were observed within 6 hours after turning 90°. (A-E) Thalli were grown under 10 days light and 9 days dark vertically (direction of the gravity 1^st^ in B), and then they were rotated at 90° (direction of the gravity 2^nd^ in B). (A) Representative images of the subapical region (500-700 um from the tip) were shown. After 3 h of 90° rotation, sedimentation of the amyloplasts started to be observed, despite the accumulation of amyloplast on the bottom of the cell in the original direction of gravity. After 6 h of 90° rotation, cells with sedimented amyloplasts increased. After 24 h, amyloplast sedimentation was clearly observed. (B-E) Statistical analysis of amyloplast distribution in cells within 700 μm from the tip. A cell was separated into four regions (I–IV) along the 2^nd^ gravity vector (bottom in B), and the area of amyloplasts in each region was counted and shown as amyloplast occupancy (%) (2^nd^ in C-E). A cell was separated into four regions (V-VIII) along the 1^st^ gravity vector (top in B), and the area of amyloplasts in each region was counted and shown as amyloplast occupancy (%) (1^st^ in C-E). Each bar indicates the average ± SD (N=8). Significant differences were identified by the Kruskal-Wallis rank sum test followed by Steel-Dwass, and are indicated with * P<0.05, **P<0.01, ***P<0.001.

### Generation of starchless mutants in M. polymorpha

To verify whether the settling of amyloplasts is necessary for sensing gravity in *M. polymorpha*, we generated mutants lacking amyloplasts. Representative starchless mutants *pgm* and *aps1* in Arabidopsis, which are deficient in plastid phosphoglucomutase and a small subunit of ADP-glucose pyrophosphorylase, required for the full development of amyloplasts, have reduced gravitropic response in both roots and shoots (Caspar and Pickard, 1989; Kiss et al., 1989, 1996; Weise and Kiss, 1999; Vitha et al., 2000). Therefore, we first identified homologs of these genes in *M. polymorpha*. Arabidopsis has three *PGM* genes (*AtPGM1-3*). Only AtPGM1 is localized in the plastid and required for starch biosynthesis (Caspar *et al*., 1985). In the *M. polymorpha* genome, we identified two homologs of *PGM* genes (Fig. 5A). The phylogenetic analysis suggested that MpPGM1 is a counterpart of AtPGM1, and MpPGM2 is that of AtPGM2/3 (Fig. 5A). Indeed, MpPGM1 but not MpPGM2 possesses the transit peptide, and Citrine-tagged MpPGM1 (MpPGM1-Citrine) was localized in the plastid, as our previous observation (Norizuki et al., 2023; Fig. 5B, C). Therefore, MpPGM1 could be involved in starch biosynthesis in plastids. ADP-glucose pyrophosphorylase is a heterotetramer (two large subunits and two small subunits) and Arabidopsis possesses two small subunit genes (At*APS1*, At*APS2*) and four large subunit genes (At*APL1-4*) (Villand *et al*., 1993). In the *M. polymorpha* genome, one small subunit gene (Mp*APS1*) and three large subunit genes (Mp*APL1-3*) were detected (Fig. 5A). MpAPS1 possesses the transit peptide and Citrine-tagged MpAPS1 was localized in the plastid (Fig. 5B, D). Taken together, we chose Mp*PGM1* and Mp*APS1* genes to generate starchless *M. polymorpha* mutant lines.

**Fig. 5.**
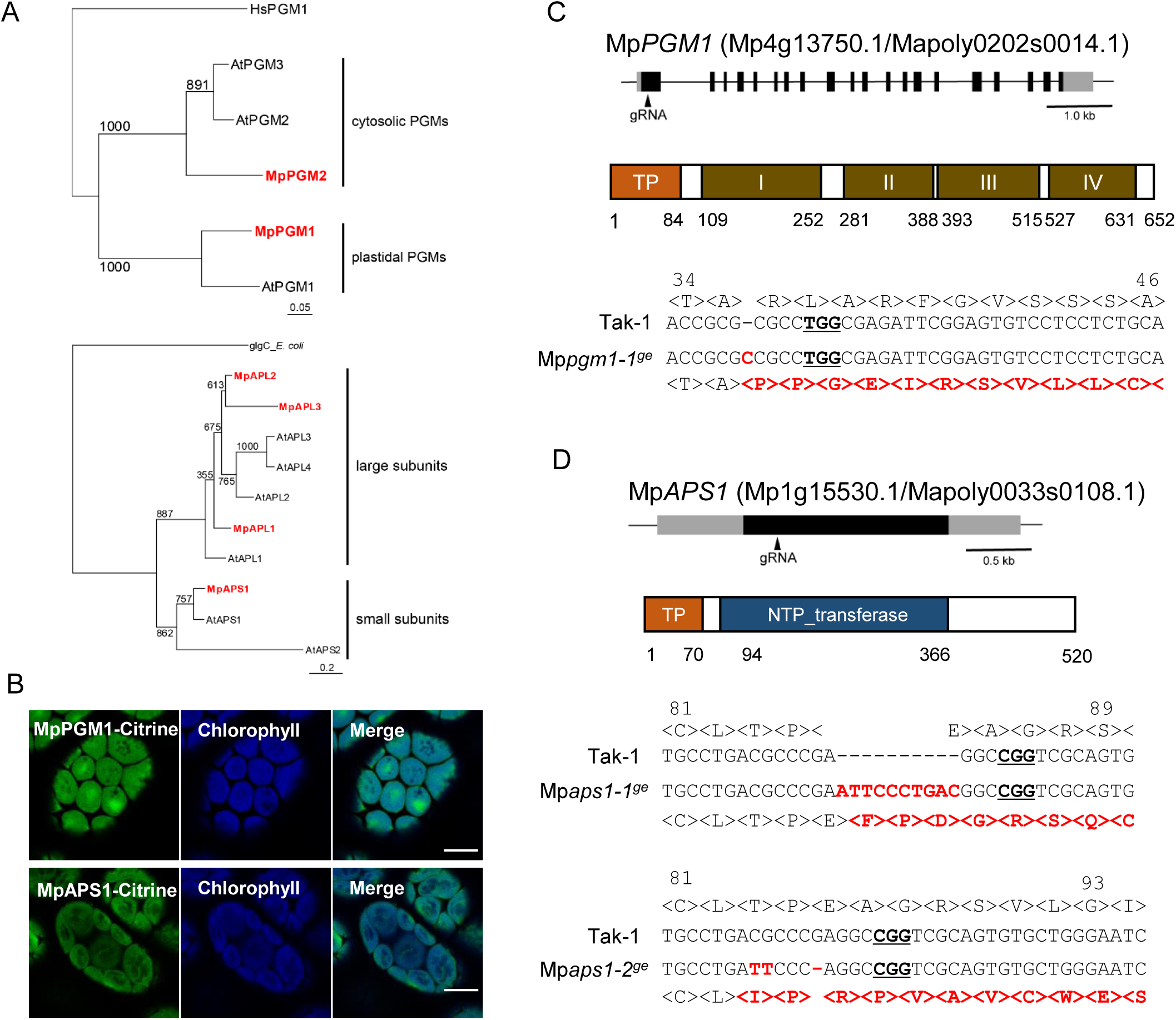
Starchless mutants in *M. polymorpha* Tak-1 produced by CRISPR-Cas9 method. (A) The maximum-likelihood phylogenetic tree of PGM and APS/APL. The maximum-likelihood phylogenetic analysis was performed using sequences of PGM, APS, and APL in Arabidopsis and *M. polymorpha*. As an outgroup to these, *Homo sapiens* PGM (for PGM) or *Escherichia coli* glgC (for APS/APL) was used. Bootstrap = 1000. (B) Localization of MpPGM1 (upper) and MpAPS1 (bottom). Green and blue pseudo-colors indicated fluorescence from Citrine and chlorophyll, respectively. Scale bars = 10 μm. (C) Mp*PGM1* gene structure and mutation site in the *Mppgm1-1* mutant. Gray and black boxes indicate untranslated and coding regions, respectively, and arrowheads indicate the region of the used guide RNA (gRNA) (upper). Domain sequences of MpPGM1. MpPGM1 possesses the transit peptide at the N terminus and the PGM_PMM_I (PF02878), PGM_PMM_II (PF02879), PGM_PMM_III (PF02880), and PGM_PMM_IV (PF00408) domains (middle). The genome and translated amino acid sequences of Mp*PGM1* genes in Tak-1 (Wild-type) and Mp*pgm1-1^ge^* (bottom). The PAM sequence for gRNA is underlined, and red letters indicate the mutation sites in Mp*pgm1-1^ge^*. (D) Mp*APS1* gene structure and mutation sites in the *Mpaps1-1* and *Mpaps1-2* mutants. Gray and black boxes indicate untranslated and coding regions, respectively, and arrowheads indicate the region of (gRNA) (upper). Domain sequences of Mp*APS1* (middle). Mp*APS1* possesses the transit peptide at the N terminus and the NTP_transferase (PF00483) domain. The genome and translated amino acid sequences of MpAPS1 genes in Tak-1, Mp*aps1-1^ge^,* and Mp*aps1-2^ge^* (bottom). The PAM sequence for gRNA is underlined, and red letters indicate the mutation sites in Mp*aps1-1^ge^* and Mp*aps1-2^ge^*.

Using CRISPR/Cas9 system (Sugano *et al*., 2018), we obtained one Mp*pgm1* and two Mp*aps1* mutant lines with frame-shift mutations (Mp*pgm1-1*^ge^, Mp*aps1-1*^ge^ and Mp*aps1-2*^ge^; Fig. 5C, D). We first examined when and where Lugol’s staining was detected in wild-type plants. In the wild type, the whole plant body was strongly stained in gemmalings on days 0 and 1, but became speckled on days 2, and 3 (Supplementary Fig. S4). The Lugol stain was limited in the center and notch areas after day 4 (Supplementary Fig. S4). We checked Lugol’s staining in gemmae and narrow structures from the wild-type Tak-1, Mp*pgm1-1^ge^*, Mp*aps1-1^ge^*, and Mp*aps1-2^ge^*. Blue-purple coloring in the wild-type Tak-1, but not in Mp*pgm1-1^ge^*, Mp*aps1-1^ge^*, and Mp*aps1-2^ge^* (Fig. 6A-H). Further detailed examination by mPS-PI showed that the signals were observed both on the surface and at a depth of 40 µm of the wild-type narrow structures (Fig. 6I, M), whereas starch grains were not detected in Mp*pgm1* and Mp*aps1* mutants (Fig. 6J-L, N-P), indicating that these mutants are starchless and do not have sedimentable amyloplasts.

**Fig. 6.**
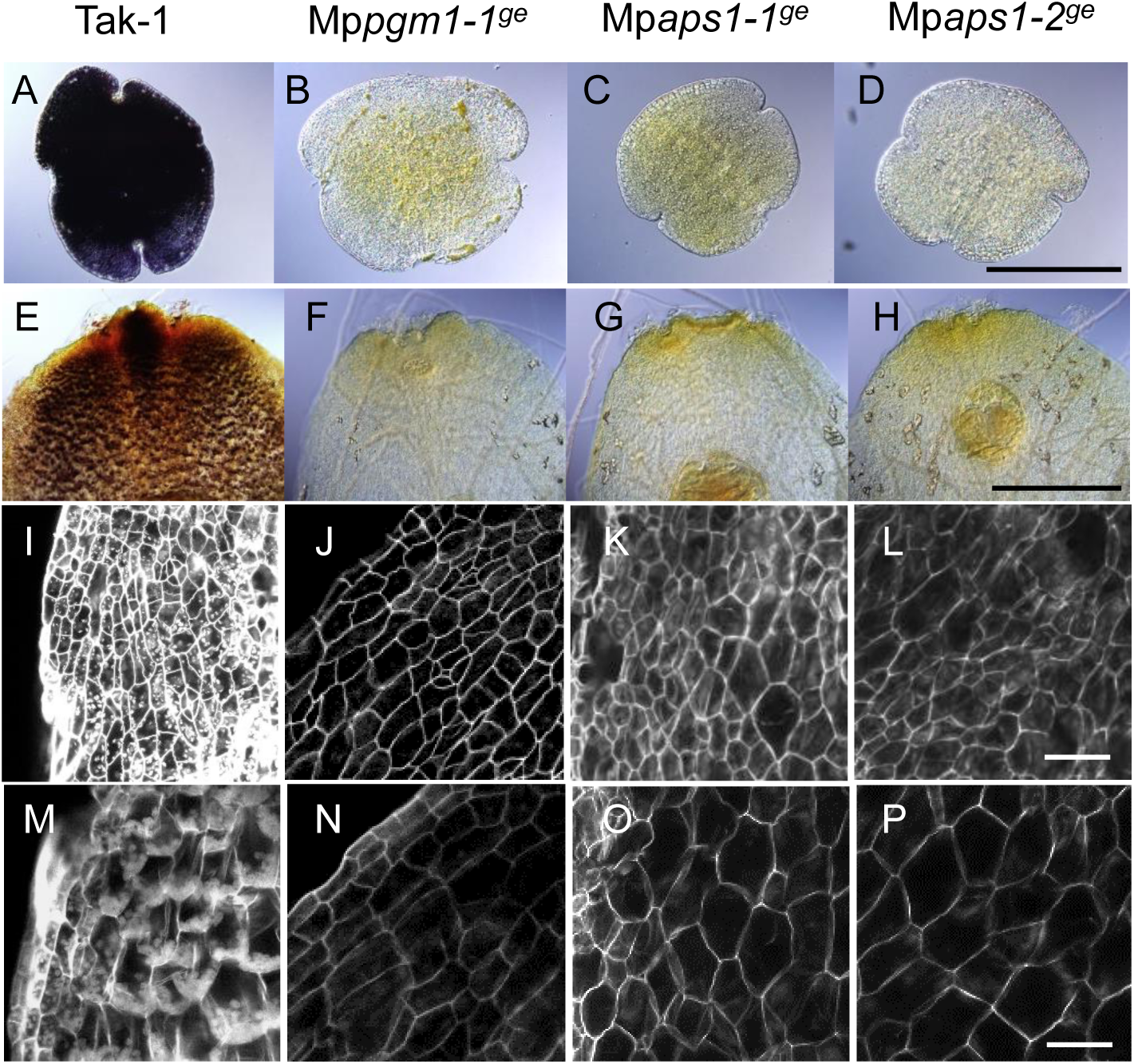
Starch stain in the wild-type plant, Mp*pgm1-1^ge^*, Mp*aps1-1^ge^*, and Mp*aps1-2^ge^* mutants. (A-H) Starch of gemmae (A-D) or narrow structures (E-H) were stained with Lugol’s solution. The narrow structures of 11-day light and 6-day vertically dark-grown thalli were used for the stain. (I-P) Narrow structures stained with modified propidium iodide staining. Optical sections of surface (I-L) and 40 µm inside (M-P) from the surface of the narrow structures. Scale bars = 200 µm in A-D; 100 µm in E-H; 50 µm in I-P.

### Starch-containing amyloplasts were required for full gravitropism in M. polymorpha

Wild-type, Mp*pgm1*, and Mp*aps1* were grown for 2 weeks in light and grown vertically for 2 weeks in dark. Narrow structures were formed, but unlike the wild-type plant, those of each mutant were not straight but helical and bent (Fig. 7A-C). To quantify the degrees of bending, the tortuosity, the length of the narrow thallus (L)/the shortest distance from the base to the top of the narrow thallus (L0), was evaluated. The value of tortuosity was almost 1 in the wild-type indicating straight growth, but was larger in Mp*pgm1-1^ge^*, Mp*aps1-1^ge^,* and Mp*aps1-2^ge^* (Fig. 7G). The length of narrow structures in the wild-type plants, Mp*pgm1-1^ge^*, Mp*aps1-1^ge^*, and Mp*aps1-2^ge^* were not significantly different (Supplementary Table S3) indicating that the narrow structures of the starchless mutants tended to be frizzier than those of the wild-type. The angles from the base to the top of the narrow structures were mostly opposed to the direction of gravity in the wild-type plants (Fig. 7H). The starchless mutants also had the highest frequency of growth in the direction opposite to that of gravity, but larger σ values indicate greater variation in the growth angle (Fig. 7H).

**Fig. 7.**
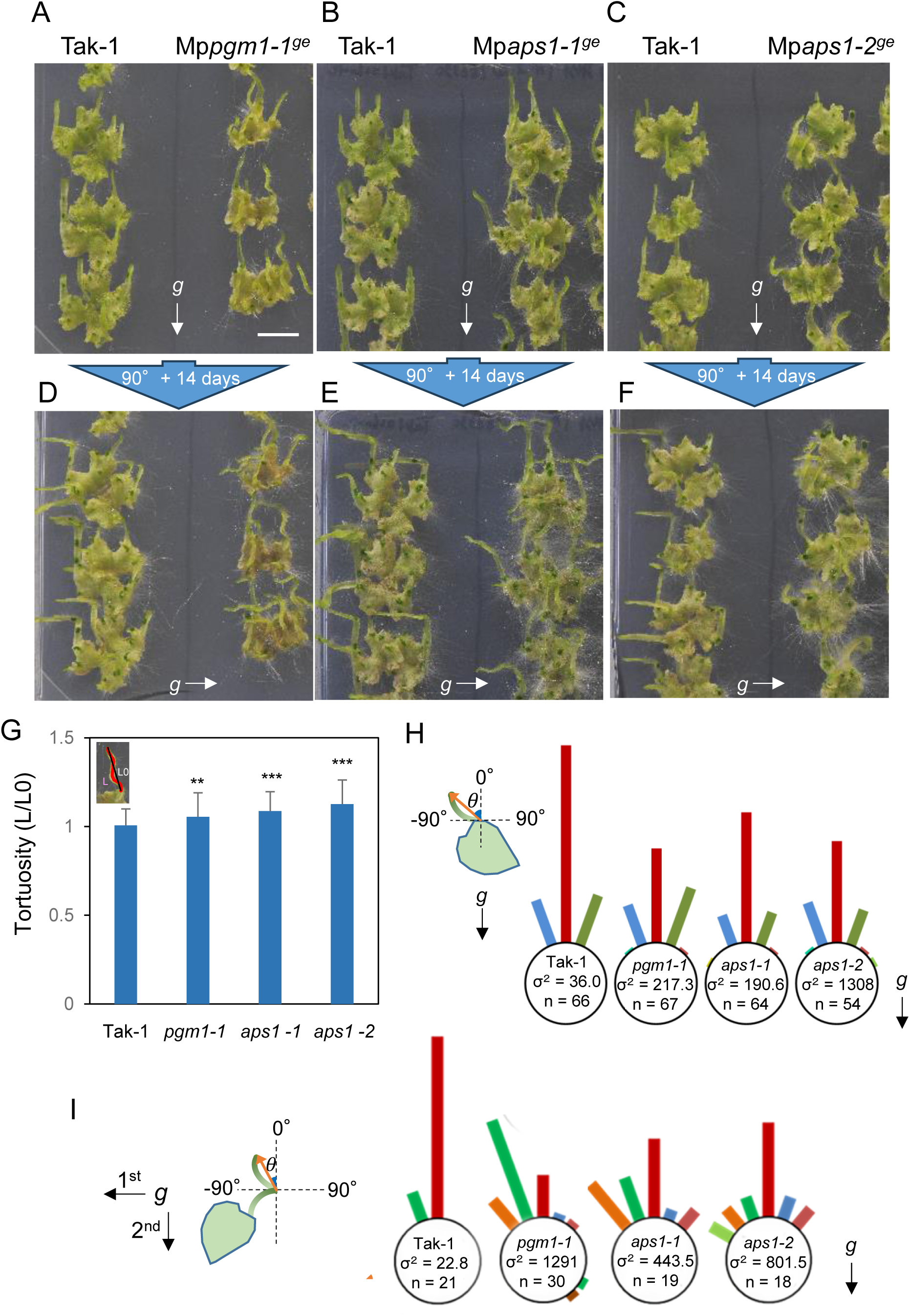
Narrow structures of starchless mutants were wavy, with greater variation in the direction of elongation. Wild-type, Mp*pgm1-1^ge^*, Mp*aps1-1^ge^,* and Mp*aps1-2^ge^* were grown in the light for 2 weeks and grown vertically in the dark for 2 weeks (A, B, and C, respectively), then they were turned 90° and grown vertically for 14 days in darkness (D, E, and F, respectively). The left lanes of each plate were Tak-1 used as controls. (G) Tortuosity of narrow structures in the wild-type plant and starchless mutants. Tortuosity is determined by the ratio of the actual length (L; red bar) to the straight distance from the basal position of the narrow structures to the tip (L0; black bar). Kruskal-Wallis rank sum test followed by Steel-Dwass, **P<0.01 or ***P<0.001, N > 54, Mean ± SD. (H) Measuring the angle (0) from the basal position of the narrow structures to the top. The bar length is proportional to the number of plants observed in each 20° bin. (I) Measuring the angle (0) from the tip of the narrow thallus before the 90° gravistimulation to the new tip of that 14 days after gravistimulation. Scale bar = 1 cm.

The gravitropic response of narrow structures was further examined when plants were rotated by 90° (Fig. 7D-F). The narrow structures of the wild-type elongated in the direction opposite to that of gravity ranging from −15.8 to 2.1°, whereas the starchless mutants elongated in various directions ranging from −44.0 to 143.7°, −43.0 to 27.0° and −54.5 to 45° in Mp*pgm1-1^ge^*, Mp*aps1-1^ge^* and Mp*aps1-2^ge^*, respectively (Fig. 7I). Reorientation induced much higher degree of variability of the growth direction of narrow structures in the starchless mutants were observed (Fig. 7I).

The reduced gravitropic phenotype of the starchless mutants may be most pronounced when the direction of gravity was changed, so we captured the dynamics of the starchless mutant when subjected to a 90-degree gravitational stimulus (Fig. 8, Supplementary Movie S3). The movements of the apical top of the narrow structures were quantitatively analyzed by single-particle tracking (Fig. 8). In the case of wild-type plants, the narrow structures elongated to the top with very slight side-to-side movements, but the swing was mostly within 1 mm to the left or right. On the other hand, starchless mutants moved more widely, moreover, many of them moved in large arcs and/or sometimes grew in the direction of gravity showing altered gravitropism (Fig. 8, Supplementary Movie S3). These results demonstrated that starch-containing amyloplasts were required for full gravitropism of thallus in *M. polymorpha* as with vascular plants such as *A. thaliana*.

**Fig. 8.**
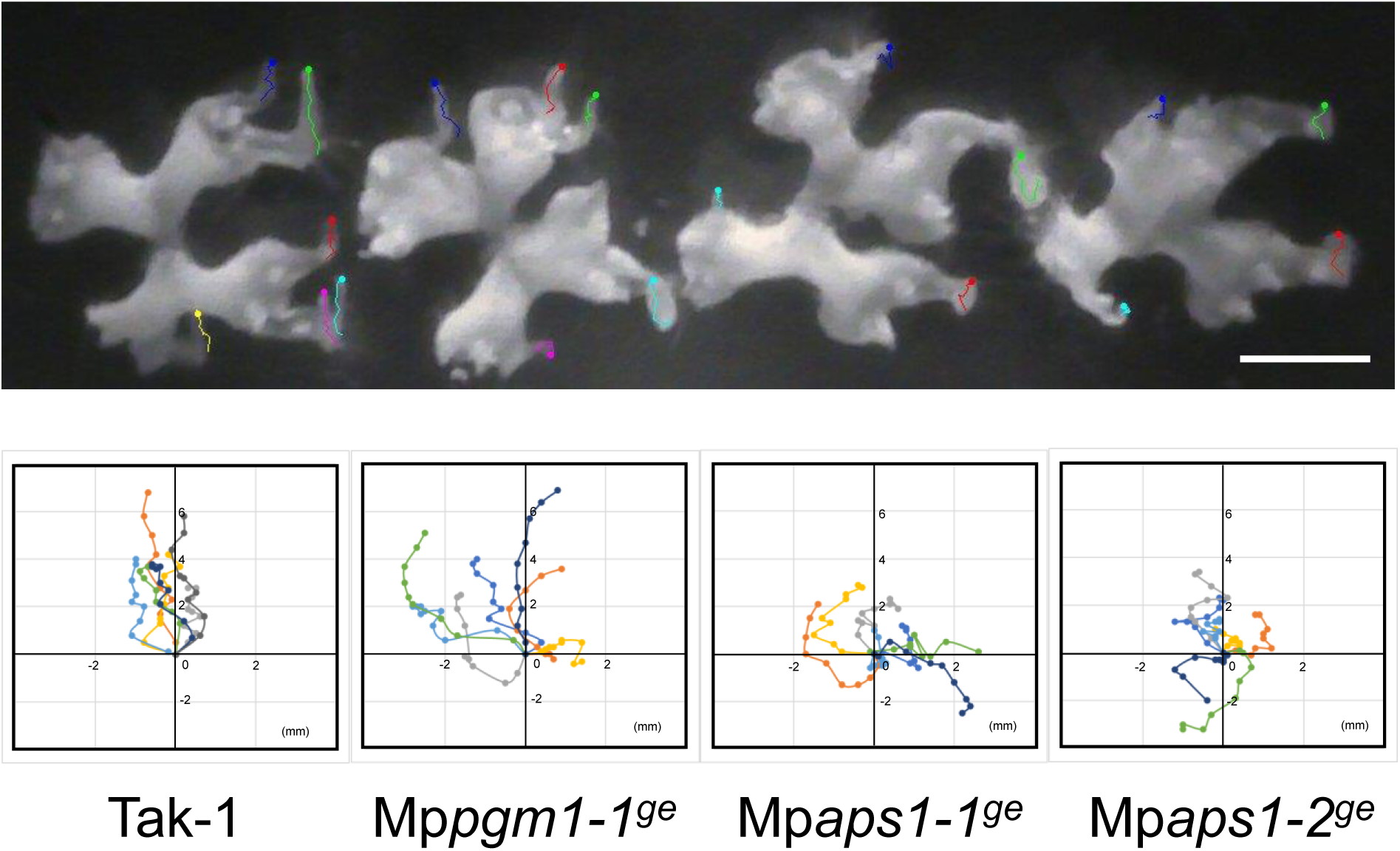
Tip movement traces in the narrow structures of the wild-type plant and starchless mutants (Mp*pgm1-1^ge^*, Mp*aps1-1^ge^*, and Mp*aps1-2^ge^*) after 90° gravistimulation. Thallus was grown for 14 days in the light and 12 days in the dark vertically and then rotated at 90°. Plots were taken every 12 hours for 5 days. N=7. Scale bar = 1 cm. The graphs show the movement in a 2D direction when the starting point is 0 ((x, y) = (0, 0))

## Discussion

### Narrow structures protruding from thalli are susceptible to gravity in M. polymorpha

We have found that structures extended from the thalli of *M. polymorpha* grew upward when they were subjected to dark conditions (Fig. 1). This direction of elongation was always opposite to gravity in response to changes in gravity directions, but 3D clinostat treatment disrupted the gravity response, resulting in the narrow structures wiggling and curving in various directions (Fig. 2). These results demonstrated that the narrow structures were highly sensitive to gravity. The plant materials commonly used to study gravitropism were cylinder-like structures, such as roots, stems, hypocotyls, and protonemata, which are susceptible to bending in any direction. Such shapes may be suitable structures for quickly determining the direction of growth in the absence of light information. The thallus of *M. polymorha* is usually flat and the rhizoids growing from them adhere to the substrate making it difficult to move. Narrow structures are one of the suitable structures for achieving sharp movements in response to gravity.

When thalli were placed in an environment without light, the original thalli stopped growing and faded in color in the dark, whereas the narrow structures were green and actively elongating (Fig. 1I), which may be due to the supply of energy and nutrients from the original thalli that accompanies the formation of the actively dividing parts (narrow structures). The narrow structures have almost the same properties as the general thalli except for a narrow body with one-directional elongation, and shallow or barely visible rims of the gemma cups on the narrow structures (Supplementary Fig. S2). Auxin is one of the factors inducing the elongation of rims of gemma cups (Kato et al. 2015), so the shallow gemma cups on narrow thallus may result from the lower level of auxin or lower sensitivity to that. Excision of the apex fully provoked regeneration and a transient decrease in endogenous auxin is involved in initiating cell cycle re-entry during regeneration (Ishida *et al*., 2022). Darkness may have a similar effect to regeneration after excitation, causing a reduction in auxin levels and thus increasing growth activity at the apical tip.

### Presence of amyloplasts in specific tissues and their sedimentation in gravity direction

It has been reported that one-day dark-treated *M. polymorpha* thalli collected from the field were treated with Lugol’s staining, and the staining appeared to be distributed unilaterally within the cells of the ground tissue including whole thalli and gametophytes (Němec, 1900; Petschow, 1933). We also detected sedimented amyloplasts in narrow structures by Lugol’s staining, but no signal was found in the original thalli placed in the dark for 2 weeks (Supplementary Fig. S1). We also showed that gemmae obtained from gemma cups were stained well overall, but the color faded after 2 days, and nothing but apical notches were stained after 4 days (Supplementary Fig. S4). These results show that starch granules are not present at any time, but only at certain times and places. These parts stained well are the critical place for gravity perception to determine the dorsoventral polarity (e.g. gemmae) or growth direction (e.g. apical notches) (Fitting, 1936; Halbsguth, 1953), which are consistent with the presence of amyloplasts as statoliths. The Lugol’s staining in narrow structures appeared to be biased toward the bottom of the cell, this pattern was very similar to that reported in the statocyte endodermal cells of Arabidopsis flower stalks, also suggesting a function of starch granules as potential statoliths (Supplementary Fig. S1, Weise and Kiss, 1999). mPS-PI method also detected amyloplasts settled in the direction of the gravity vector in parenchyma cells of the narrow structures (Fig. 3). This method also detected a signal of small particles in the epidermis, which did not settle intracellularly, suggesting that these were plastids with insufficient accumulation of starch (Fig. 3).

Sedimentation of amyloplasts was recognized before the morphological changes caused by gravistimulation of the narrow structures. The fact that amyloplast sedimentation preceded gravitropic bending is consistent with the starch-statolith hypothesis. It has been reported that sedimentation of amyloplasts in shoot endodermal cells of Arabidopsis observed within 3 minutes at 90° reorientation (Saito et al., 2005), and Arabidopsis flowering stalks achieved 60°curvature in 80 min (Caspar and Pickard, 1989). These time courses seem considerably faster than those were demonstrated at narrow structures of *M. polymorpha* (Figs 2 and 4). In roots or rhizoids, gravitropic bending in seed plants after a 90° reorientation of the seedlings is faster than that of basal vascular plants including ferns and lycophytes, and non-vascular plants such as mosses (Zhang *et al*., 2019). Similar to the root system, gravitropism of shoots or thalli may have developed with evolution through restricted statocytes and their structural changes to facilitate the settling of statoliths.

### Amyloplasts are statoliths in M. polymorpha as in seed plants

Gravity-sensing mechanisms differ among organisms. Unicellular green algae, *Chlamydomonas reinhardtii* sense gravity using the whole body rather than starch-covered pyrenoid (Roberts, 2006), and *Chara globularis* using barium sulfate vesicles rather than starch-filled amyloplasts (Limbach *et al*., 2005). In contrast to green algae, the protonemata or caulonemata of mosses possess sedimentable starch-filled amyloplasts in their growing tips, whose role as the statolith is suggested (Jenkinos et al., 1986; Walker and Sack, 1990; Schwuchow et al., 1995; Sack et al., 2001). To test whether amyloplast sedimentation contributes to gravity sensing in *M. polymorpha*, we generated starch-deficient Mp*pgm1* and Mp*aps1* mutants with no starch signals (Figs. 5, 6). Different from a straight vertical growth of narrow structures in the wild-type plant, those of the starchless mutants showed curve or spiral growth to the upper direction but with greater variability and significantly higher tortuosity (Fig. 7A-C, G, H). These results are similar to what was observed in the starchless mutants of Arabidopsis *pgm1* and *aps1*, in which vertical elongation in shoot and root is disturbed, and the variation in their growth direction is greater than that of the wild-type plant (Vitha et al., 2000). Many terrestrial plants examined so far use starch grains (amyloplasts) for gravitropism, suggesting their common role as statoliths for gravity perception in land plants (Barlow, 1995; Sievers et al., 1996; Sack et al., 2001; Yoder et al., 2001). Sufficient evidence for the role of the amyloplasts as statoliths has been obtained by using intracellular magnetophoresis to move them laterally in various plant organs and induce a gravitropic response (Kuznetsov and Hasenstein, 1997; Weise et al., 2000). The alteration of growth direction in the narrow structures of starchless mutants was more pronounced when the direction of gravity was changed (Fig. 7I), especially within 24 h of changing the direction of gravity (Fig. 8), suggesting that the gravitropism in *M. polymorpha* requires sedimented amyloplasts.

However, starchless mutants including Mp*pgm1* and Mp*aps1* can respond to gravity to some extent, and they eventually move upward after a sufficient time after gravistimulation, suggesting that there may be an amyloplast-independent pathway for the gravitropism in *M. polymorpha*. This observation is different from clinostat-treated plants. It has been observed and discussed from studies with starchless mutants of Arabidopsis that gravitropism is not completely eliminated in the absence of starch (Caspar and Pickard, 1989; Kiss et al., 1989, 1996; Kiss, 2000; Morita, 2010). Plastids not accumulating starch granules might be a reasonable candidate for statolith albeit the function is very weak, assuming that rapid and extensive plastid sedimentation is not necessary for gravity sensing. A candidate of the non-statolith system is the mass of the entire protoplast. This ‘protoplast pressure hypothesis’ is a mechanism that senses pressure due to the weight of the cytoplasm acting on the membrane and cell wall, and is thought to be an early evolved and probably less sensitive one (Wayne *et al*. 1990; Weise and Kiss, 1999). Multiple pathways may co-exist in gravitropism in land plants (Barlow, 1995).

Further studies must be needed to elucidate mechanisms of gravitropism in each plant and why the sedimentation speed of amyloplasts differs among species; how the intracellular structure evolves to facilitate gravity sensing. Comparative gravitropic studies using a wide range of experimental model plants would help to elucidate the basic mechanism acquired by the ancestral terrestrial plants and their diversification during land plant evolution.

## Abbreviations

APS: small subunit of ADP-glucose pyrophosphorylase
PGM,: plastidial phosphoglucomutase

## Supplementary data

**Fig. S1**

Amyloplasts on the narrow structures were stained with Lugol’s solution

**Fig. S2**

Narrow structures retained some traits as thallus

**Fig. S3**

Narrow structures came out from the thalli buried in the ground

**Fig. S4**

Young gemmalings lose their starch accumulation as they grow

**Table S1**

Primers used in this study

**Table S2**

The length of narrow structures treated with and without clinorotation

**Table S3**

The length of narrow structures in WT and starchless mutants

**Material S1**

The sequences used in the phylogenetic analysis of PGM

**Material S2**

The sequences used in the phylogenetic analysis of APL

**Movie S1**

Time-lapse images of 11-day-old light-grown thalli transferred to the dark

**Movie S2**

Time-lapse images of narrow structures growing after 90° rotation in darkness

**Movie S3**

Time-lapse movie of narrow structures of wild-type, Mp*pgm1-1^ge^*, Mp*aps1-1^ge^*, and Mp*aps1-2^ge^* with tip traces as shown in Fig. 8

## Acknowledgements

We thank Aino Komatsu (Tohoku University, Japan), Ryuichi Nishihama (Tokyo University of Science, Japan), and Takayuki Kohch (Kyoto University, Japan) for helpful advice on *M. polymorpha*. Miya Mizutani (Nagoya University, Japan) for valuable advice on the mPS-PI method, and Hisae Kojima, Yuri Komada, and Eriko Sawada for technical assistance. We also thank Model Plant Section, Model Organisms Facility, NIBB Trans-Scale Biology Center for providing plant cultivation facilities.

## Author Contributions

MHS and MMT designed the experiments. MHS wrote and finalized the manuscript with TN, NS, and MMT. MHS analyzed the gravitropism of narrow structures. TN identified the MpPGM1 and MpAPS1 genes and established their knockout mutant lines. MHS and TN performed microscopic analysis. MHS, SS, and YO did the image analysis for quantification. The paper was supervised by TU.

## Data availability

The data underlying this article will be shared with the corresponding author, upon reasonable request.

## References

Barlow PW. 1995. Gravity perception in plants: a multiplicity of systems derived by evolution? Plant, Cell & Environment 18, 951–962.

Blancaflor EB, Masson PH. 2003. Plant Gravitropism. Unraveling the Ups and Downs of a Complex Process. Plant Physiology 133, 1677–1690.

Bowman JL. 2022. The liverwort *Marchantia polymorpha*, a model for all ages. Current Topics in Developmental Biology. 147. 1–32.

Bowman JL, Araki T, Arteaga-Vazquez MA, et al. 2016. The naming of names: Guidelines for gene nomenclature in *Marchantia*. Plant and Cell Physiology 57, 257–261.

Bowman JL, Kohchi T, Yamato KT, et al. 2017. Insights into Land Plant Evolution Garnered from the *Marchantia polymorpha* Genome. Cell 171, 287–304.e15.

Braun M, Sievers A. 1993. Centrifugation causes adaptation of microfilaments Studies on the transport of statoliths in gravity sensing Chara rhizoids. Springer-Verlag.

Caspar T, Huber SC, Somerville C. 1985. Alterations in Growth, Photosynthesis, and Respiration in a Starchless Mutant of Arabidopsis thaliana (L.) Deficient in Chloroplast Phosphoglucomutase Activity. Plant Physiology 79, 11–17.

Caspar T, Pickard BG. 1989. Gravitropism in a starchless mutant of Arabidopsis - Implications for the starch-statolith theory of gravity sensing. Planta 177, 185–197.

Emanuelsson O, Nielsen H, von Heijne G. 1999. ChloroP, a neural network-based method for predicting chloroplast transit peptides and their cleavage sites. Protein Science 8, 978–984.

Fitting H. 1936. Untersuchungen über die Induktion der Dorsiventralität bei den keimenden Brutkörpern von Marchantia und Lunularia. I. Die Induktoren und ihre Wirkungen. Jahrbücher für wissenschaftliche Botanik 82, 333–376.

Flütsch S, Distefano L, Santelia D. 2018. Quantification of Starch in Guard Cells of Arabidopsis thaliana. Bio-Protocol 8.

Gamborg OL, Miller RA, Ojima K. 1968. Nutrient requirements of suspension cultures of soybean root cells. Experimental Cell Research 50, 151–158.

Haberlandt G. 1900. Ueber die Perception des geotropischen Reizes. Berichte der Deutschen Botanischen Gesellschaft 18, 261–272.

Halbsguth W. 1953. Über die Entwicklung der Dorsiventralität bei *Marchantia polymorpha L*. Ein Wuchsstoff problem? Biologisches Zentralblatt 72, 52–104.

Hangarter RP. 1997. Gravity, light and plant form. Plant, Cell and Environment 20, 796–800.

Haseloff J. 2003. BioImaging Old Botanical Techniques for New Microscopes. Bioimaging 34, 1174–1182.

Hodick D, Buchen B, Sievers A. 1998. Statolith positioning by microfilaments in *Chara* rhizoids and protonemata. Advances in Space Research 21, 1183–1189.

Hoson T, Kamisaka S, Masuda Y, Yamashita M. 1992. Changes in plant growth processes under microgravity conditions simulated by a three-dimensional clinostat. The Botanical Magazine Tokyo 105, 53–70.

Ishida S, Suzuki H, Iwaki A, et al. 2022. Diminished Auxin Signaling Triggers Cellular Reprogramming by Inducing a Regeneration Factor in the Liverwort *Marchantia polymorpha*. Plant and Cell Physiology 63, 384–400.

Ishizaki K, Nishihama R, Ueda M, Inoue K, Ishida S, Nishimura Y, Shikanai T, Kohchi T. 2015. Development of gateway binary vector series with four different selection markers for the liverwort *Marchantia polymorpha*. PLoS ONE 10, 1–13.

Jenkins GI, Courtice GRM, Cove DJ. 1986. Gravitropic responses of wildCtype and mutant strains of the moss *Physcomitrella patens*. Plant, Cell & Environment 9, 637–644.

Kato H, Ishizaki K, Kouno M, Shirakawa M, Bowman JL, Nishihama R, Kohchi T. 2015. Auxin-Mediated Transcriptional System with a Minimal Set of Components Is Critical for Morphogenesis through the Life Cycle in *Marchantia polymorpha*. PLoS genetics. 1(5): e1005084.

Kiss JZ. 2000. Mechanisms of the Early Phases of Plant Gravitropism. Critical Reviews in Plant Sciences 19. 551–573.

Kiss JZ, Guisinger MM, Miller AJ, Stackhouse KS. 1997. Reduced Gravitropism in Hypocotyls of Starch-Deficient Mutants of Arabidopsis. Plant and Cell Physiology 38, 518– 525.

Kiss JZ, Hertel R, Sack FD. 1989. Amyloplasts are necessary for full gravitropic sensitivity in roots of Arabidopsis thaliana. Planta 177, 198–206.

Kiss JZ, Wright JB, Caspar T. 1996. Gravitropism in roots of intermediate-starch mutants of Arabidopsis. Physiologia Plantarum 97, 237–244.

Knight TA. 1806. On the direction of the radicle and germen during the vegetation of seeds. Philosophical Transactions of the Royal Society of London 96, 99–108.

Kohchi T, Yamato KT, Ishizaki K, Yamaoka S, Nishihama R. 2021. Development and Molecular Genetics of *Marchantia polymorpha*. Annual Review of Plant Biology. 72:677–702.

Konings H. 1995. Gravitropism of during the last three decades. 44, 195–223.

Kubota A, Ishizaki K, Hosaka M, Kohchi T. 2013. Efficient Agrobacterium-mediated transformation of the liverwort *Marchantia polymorpha* using regenerating thalli. Bioscience, Biotechnology and Biochemistry 77, 167–172.

Kurihara D, Mizuta Y, Sato Y, Higashiyama T. 2015. ClearSee: A rapid optical clearing reagent for whole-plant fluorescence imaging. Development 142, 4168–4179.

Kuznetsov OA, Hasenstein KH. 1997. Magnetophoretic induction of curvature in coleoptiles and hypocotyls. Journal of Experimental Botany 48, 1951–1957.

Lefort V, Longueville JE, Gascuel O. 2017. SMS: Smart Model Selection in PhyML. Molecular Biology and Evolution 34, 2422–2424.

Letunic I, Bork P. 2018. 20 years of the SMART protein domain annotation resource. Nucleic Acids Research 46, D493–D496.

Letunic I, Khedkar S, Bork P. 2021. SMART: Recent updates, new developments and status in 2020. Nucleic Acids Research 49, D458–D460.

Limbach C, Hauslage J, Schäfer C, Braun M. 2005. How to activate a plant gravireceptor. Early mechanisms of gravity sensing studied in characean rhizoids during parabolic flights. Plant Physiology 139, 1030–1040.

Montgomery SA, Tanizawa Y, Galik B, et al. 2020. Chromatin Organization in Early Land Plants Reveals an Ancestral Association between H3K27me3, Transposons, and Constitutive Heterochromatin. Current Biology 30, 573–588.e7.

Morita MT. 2010. Directional gravity sensing in gravitropism. Annual Review of Plant Biology 61, 705–720.

Morita MT, Tasaka M. 2004. Gravity sensing and signaling. Current Opinion in Plant Biology 7, 712–718.

Němec B. 1900. Über die Art der Wahrnehmung des Schwerkraftreizes bei den Pflanzen. Berichte der Deutschen Botanischen Gesellschaft 18, 241–245.

Norizuki T, Kanazawa T, Minamino N, Tsukaya H, Ueda T. 2019. Marchantia polymorpha, a new model plant for autophagy studies. Frontiers in Plant Science 10, 1–16.

Norizuki T, Minamino N, Sato M, Ueda T. 2023. Autophagy regulates plastid reorganization during spermatogenesis in the liverwort Marchantia polymorpha. Frontiers in Plant Science 14.

Petschow F. 1933. Geotropismus und Statolithenstärke bei Bryophyten. Beihefte zum Botanischen Centralblatt 51, 287–310.

Roberts AM. 2006. Mechanisms of Gravitaxis in Chlamydomonas. The biological bulletin 210, 78–80.

Saito C, Morita TM, Kato T, Tasaka M. 2005. Amyloplasts and Vacuolar Membrane Dynamics in the Living Graviperceptive Cell of the Arabidopsis Inflorescence Stem. The Plant Cell, 17, 548–558.

Sack FD. 1991. Plant Gravity Sensing. International Review of Cytology. 127, 193–252.

Sack FD, Schwuchow JM, Wagner T, Kern V. 2001. Gravity sensing in moss protonemata. Advances in Space Research 27, 871–876.

Schindelin J, Arganda-Carrera I, Frise E, et al. 2009. Fiji - an Open platform for biological image analysis. Nature Methods 9.

Schwuchow JM, Kim Donggiun, Sack FD. 1995. Caulonemal gravitropism and amyloplast sedimentation in the moss Funaria. Canadian Journal of Botany 73, 1029–1035.

Segami S, Tomoyama T, Sakamoto S, Gunji S, Fukuda M, Kinoshita S, Mitsuda N, Ferjani A, Maeshima M. 2018. Vacuolar H^+^-pyrophosphatase and cytosolic soluble pyrophosphatases cooperatively regulate pyrophosphate levels in Arabidopsis thaliana. Plant Cell 30, 1040–1061.

Shimamura M. 2016. *Marchantia polymorpha*: Taxonomy, phylogeny and morphology of a model system. Plant and Cell Physiology 57, 230–256.

Sievers A, Buchen B, Hodick D. 1996. Gravity sensing in tip-growing cells. Trends in Plant Science. 1, 273–279.

Sievers A, Kramer-Fischer M, Braun M, Buchen B. 1991. The Polar Organization of the Growing *Chara* Rhizoid and the Transport of Statoliths are Actin-Dependent. Botanica Acta 104, 103–109.

Sugano SS, Nishihama R, Shirakawa M, Takagi J, Matsuda Y, Ishida S, Shimada T, Hara-Nishimura I, Osakabe K, Kohchi T. 2018. Efficient CRISPR/Cas9-based genome editing and its application to conditional genetic analysis in Marchantia polymorpha. PLoS ONE 13, 1–22.

Tasaka M, Kato T, Fukaki H. 1999. The endodermis and shoot gravitropism. Trends in Plant Science 4, 103–107.

Villand P, Olsen O-A, Kleczkowski LA. 1993. Molecular characterization of multiple cDNA clones for ADP-glucose pyrophosphorylase from *Arabidopsis thaliana*. Plant Molecular Biology. 23. 1279–1284.

Vitha S, Zhao L, Sack FD. 2000. Interaction of root gravitropism and phototropism in Arabidopsis wild-type and starchless mutants. Plant Physiology 122, 453–461.

Walker LM, Sack FD. 1990. Amyloplasts as possible statoliths in gravitropic protonemata of the moss *Ceratodon purpureus*. Planta 181, 71–77.

Wayne R, Staves MP, Leopold AC. Gravity-dependent polarity of cytoplasmic streaming in *Nitellopsis*. Protpplasma 155, 43–57.

Weise SE, Kiss JZ. 1999. Gravitropism of inflorescence stems in starch-deficient mutants of Arabidopsis. International Journal of Plant Sciences 160, 521–527.

Weise SE, Kuznetsov OA, Hasenstein KH, Kiss JZ. 2000. Curvature in Arabidopsis inflorescence stems is limited to the region of amyloplast displacement. Plant and Cell Physiology 41, 702–709.

Yoder T, Zheng H, Todd P, Staehelin L. 2001. Amyloplast sedimentation dynamics in maize columella cells support a new model for the gravity-sensing apparatus of roots. Plant Physiology 125,1045–1060.

Zhang Y, Xiao G, Wang X, Zhang X, Friml J. 2019. Evolution of fast root gravitropism in seed plants. Nature Communications 10, 4–13.

