## Supplementary data for "Amyloplasts are necessary for full gravitropism in thallus of *Marchantia polymorpha*"

#### Amyloplasts are necessary for full gravitropic sensitivity in *M. polymorpha*

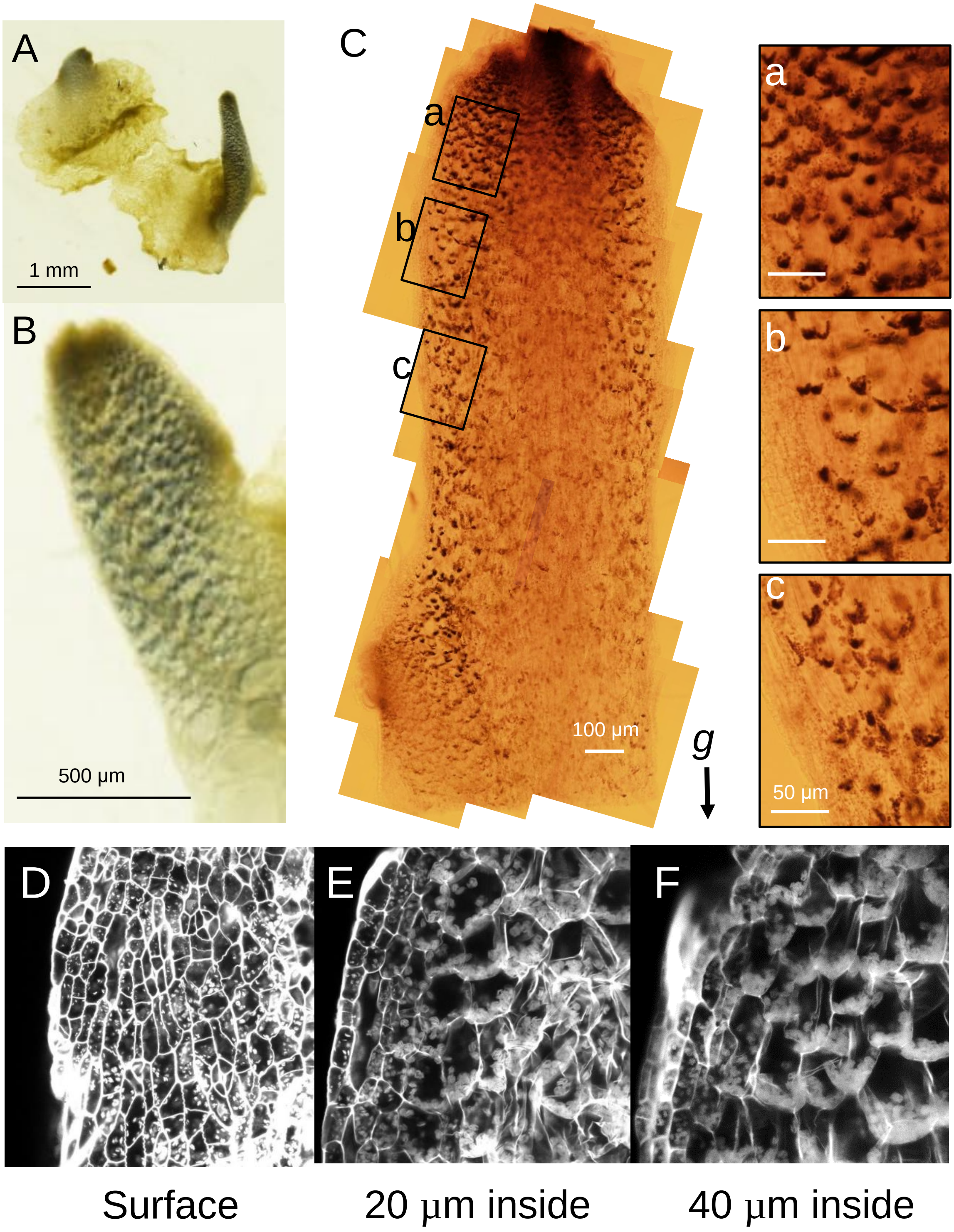


Fig. S1

Amyloplasts on the narrow structures were stained with Lugol's solution. (A, B) 7-day-old light-grown thalli were transferred to darkness and vertically grown for 14 days. Narrow strucutres come out from thalli were stained in dots or speckled but the original thalli were not. (C) Representative Lugol stained narrow structures of a thallus, grown in the light for 2 weeks and grown vertically in the dark for 2 weeks. (a-c) Magnified images of C; The regions of approximately 200 µm to 400 µm (a), 500 µm to 700 µm (b) 800 µm to 1 mm (c) from the tip. Cells closer to the tip had more stained starch granules and the signals become indistinct toward the base of the narrow structures.

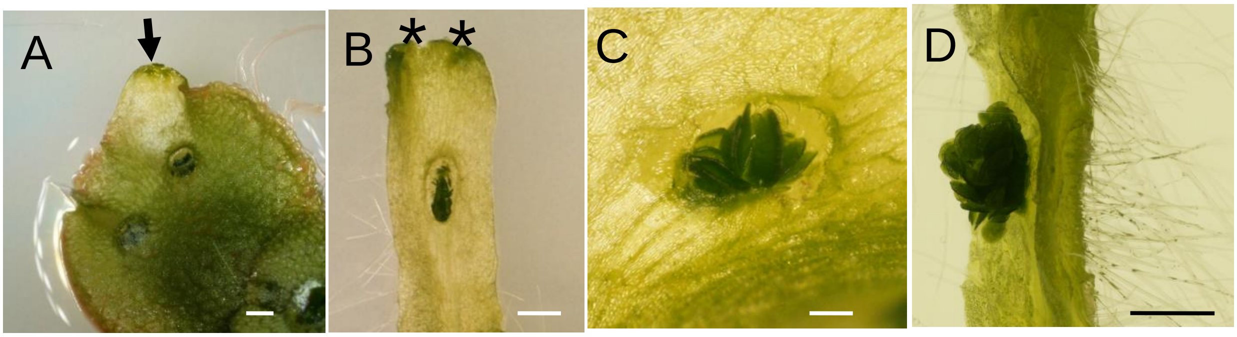


### Fig. S2

Narrow structures retained some traits as thallus. (A) Emergence of a bud structure (black arrow) from an apical notch of the thallus grown for 2 weeks in the light and 5 days in the dark. (B) Narrow structures grew straight with one or two apical notches (asterisks) at the apex. (C, D) Developed narrow structures produced some gemma cups along the midrib. Developing (C) and mature (D) gemma cups on the narrow structures with very shallow rim. (D) The narrow structures clearly showed dorsiventrality; Gemma cups formed on the dorsal side, and rhizoids covered on the ventral surface. Scale bars = 1 mm in A, B, and D; 200 µm in C.

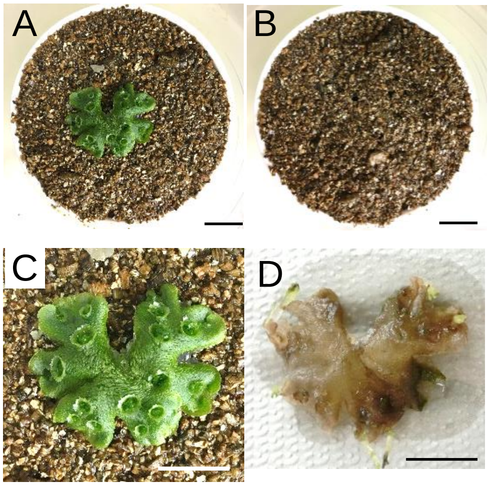


### Fig. S3

Narrow structures came out from the thalli buried in the ground. 2 week-old light-grown thalli (A) were buried in the soil at a depth of 1 cm (B). Plants before (C) and after (D) burial in the ground for 5 weeks, respectively. Scale bars = 1cm.

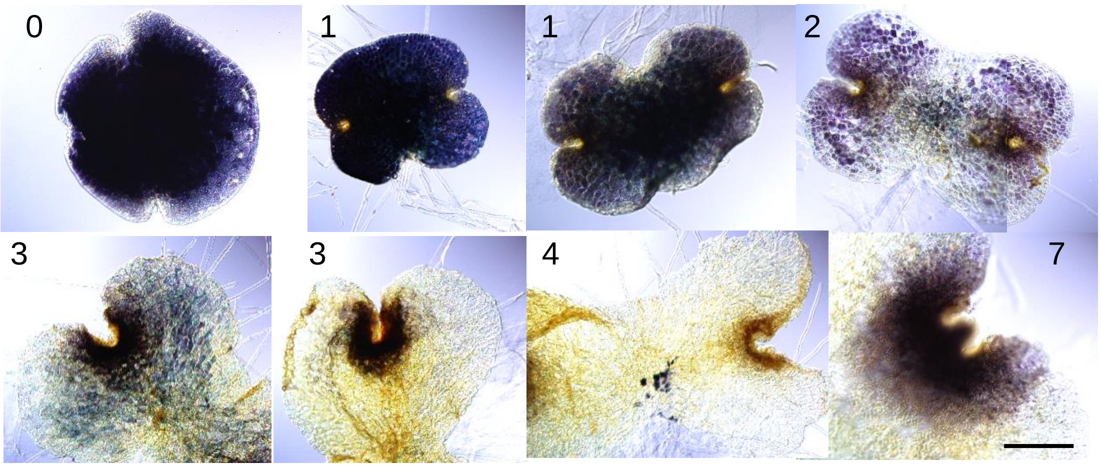


Fig. S4

Young gemmalings lose their starch accumulation as they grow. Amyloplasts in the light-grown gemmalings were stained by Lugol solution. Lugol-stained starch granules showed dark blue or purple. The numbers of the day elapsed since gemmae were shown at the top. Scale bar = 100 µm.

| Primer name | Sequences |
| --- | --- |
| MpApS1_Fw | CACCATGGCTGGTGTTGCGACAG |
| MpApS1_Rv | GATGACGGTGTCGTTGGGAA |
| MpAPS1_gDNA_Fw | CTCGAGTTGCCTGACGCCCGAGGC |
| MpAPS1_gDNA_Rv | AAACGCCTCGGGCGTCAGGCAACT |
| MpAPS1_GT1_Fw | CTTACTGCTGCTCGCTTACCT |
| MpAPS1_seq1_Fw | GCACAACCGCTGAGGTCG |
| MpPGM1_gDNA_Fw | CTCGGCATGGCGAGACCGCGCGCC |
| MpPGM1_gDNA_Rv | AAACGGCGCGCGGTCTCGCCATGC |
| MpPGM1_GT1_Fw | AAGTGGACCGCCCACGGCAT |
| MpPGM1_GT2_Rv | CCACTAGTTCCAGTCTTCTGGC |
| MpPGM1_seq1_Fw | GCGCGCACAGATGCGAAT |

Table S1 Primers used in this study

| | The length of narrow thalli (Mean ± SE cm)) |
|---|---|
| Tak-1 (1G) | 1.13 ± 0.06 |
| Clinostat | 1.20 ± 0.05 |

No significant difference (N=58, Student's t-test P=0.069)

### Table S2

The length of narrow structures treated with and without clinorotation

| | The length of narrow thalli (Mean ± SE cm) |
|---|---|
| Tak-1 | 0.80 ± 0.09 |
| Mp*pgm1-1*$^{ge}$ | 0.78 ± 0.05 |
| Mp*aps1-1*$^{ge}$ | 0.96 ± 0.05 |
| Mp*aps1-2*$^{ge}$ | 0.73 ± 0.07 |

No significant differences (N=24, One-way ANOVA, P=0.08).

### Table S3

The length of narrow structures in WT and starchless mutants

>AtPGM3
TSPIDGQKPG TSGLRKKVKV FNYLENFVQA TFNALTAEKV KGATLVVSGD GRYYSKDAVQ
IIIKMAAANG VRRVWVGKNT LLSTPAVSAV IRERKATGAF ILTASHNPGG PTEDFGIKYN
MENGGPAPES ITDKIYENTK TIKEYPIAPN VDISAVGVTS FGKFDVEVFD PADDYVKLMK
SIFDFEAIRK LLSSPFTFCY DALHGVAGAY AHRIFVEELG AQESALLNCT PKEDFGGGHP
DPNLTYAKEL VARMPEFGAA ADGDADRNMI LGKFFVTPSD SVAIIAANAA IPYFSSGLKG
VARSMPTSAA LDVVAKSLNL KFFEVPTGWK FFGNLMDAGM CSVCGEESFG TGSDHIREKD
GIWAVLAWMS ILAHKNVEDI VRQHWATYGR HYYTRYDYEN VDAGKAKELM EHLVKLQSVA
SVASADEFEY KDPVDGSISK HQGIRYLFED GSRLVFRLSG TGSEGATIRL YIEQYEKDAS
KTGRESQEAL SPLVDLALKL SKMEEFTGRS APTVIT
>MpPGM2
TKPFDGQKPG TSGLRKKVKV FHYLANFVQS SFNALPADRV KGSTIVVSGD GRYYSKDAIQ
IIIRIAAANG VAQVWVGQDG LLSTPAVSGI IRDRKAYGGF ILTASHNAGG PDEDFGIKYN
TENGGPALEA LTDQIYENTK TITTYLIAPD IDIHKIATSS FGPFSVEVFD ATEDYVKMMK
QIFDFEAISR LLARPFTFCY DALHGVAGVY AKRIFLQELG AQESSLLNCT PKEDFGGGHP
DPNLTYAKQL VKTMPEFGAA ADGDADRNMI LGKFFVTPSD SVAIIAANAA IPYFRNGLKG
VARSMPTSAA LDVVAKGLGL RQFEVPTGWK FFGNLMDAGM CSICGEESFG TGSDHIREKD
GIWAVLAWLS ILAYRNVEDI VTLHWGIYGR HYYTRYDYEN VDAGAAKNLM SHLVKLQSVS
GVKSGDEFEY KDPVDGSVSS HQGIRFLFKD GSRLVFRLSG TGSVGATIRL YIEQYQSDKS
KTGADASETL APLVDVALVL SKMEEFTGRS APTVIT
>MpPGM1
TTPIDGQKTG TSGLRKKVKE FNYLANWIQA LFDSLPAEDV KGSTLVLGGD GRYFNKEASQ
IIIKIAAGNG VGKILVGREG IASTPAVSAI IRARKANGGF VMSASHNPGG PKYDWGIKFN
YSSGQPAPES ITDKIYGNTL SIKEIKQAPD VNLSELGVHK FGDFSVEVID PVADYLNLLE
EVFDFDLLKG LLTSKFRFKF DAMHAVTGAY AKPIFVDRLG APEDSIFNGV PLEDFGGGHP
DPNLTYAEEL VKIMPDFGAA SDGDGDRNMI LGNFFITPSD SVAMIAANAA IPYFKTGLKG
LARSMPTSGA LDRVAKELGL PFFETPTGWK FFGNLMDAGK CSVCGEESFG TGSDHVREKD
GIWAVLAWIS IVAYKNVADI AKEHWAKYGR NFFSRYDYEE CESAGANKMV EHLRDIIANY
ELELADDFAY TDPIDGSVAT KQGIRFIFSD GSRIIFRLSG TGSAGATIRI YVEQYEQDTT
KHDLDAQDAL KPLIDIALSV SKLQEFTGRT KPTVIT
>AtPGM2
TSPIDGQKPG TSGLRKKVKV FNYLENFVQA TFNALTTEKV KGATLVVSGD GRYYSEQAIQ
IIVKMAAANG VRRVWVGQNS LLSTPAVSAI IRERKATGAF ILTASHNPGG PTEDFGIKYN
MENGGPAPES ITDKIYENTK TIKEYPIAPR VDISTIGITS FGKFDVEVFD SADDYVKLMK
SIFDFESIKK LLSYPFTFCY DALHGVAGAY AHRIFVEELG APESSLLNCV PKEDFGGGHP
DPNLTYAKEL VARMPEFGAA ADGDADRNMI LGKFFVTPSD SVAIIAANAA IPYFSSGLKG
VARSMPTSAA LDVVAKNLGL KFFEVPTGWK FFGNLMDAGM CSVCGEESFG TGSDHIREKD
GIWAVLAWLS ILAHKNVEDI VRQHWATYGR HYYTRYDYEN VDATAAKELM GLLVKLQSVA
NVASADEFEY KDPVDGSVSK HQGIRYLFED GSRLVFRLSG TGSEGATIRL YIEQYEKDAS
KIGRDSQDAL GPLVDVALKL SKMQEFTGRS SPTVIT
>AtPGM1
TKPIEGQKTG TSGLRKKVKV FNYLANWIQA LFNSLPLEDY KNATLVLGGD GRYFNKEASQ
IIIKIAAGNG VGQILVGKEG ILSTPAVSAV IRKRKANGGF IMSASHNPGG PEYDWGIKFN
YSSGQPAPET ITDKIYGNTL SISEIKVAPD IDLSQVGVTK YGNFSVEVID PVSDYLELME
DVFDFDLIRG LLSRSFGFMF DAMHAVTGAY AKPIFVDNLG AKPDSISNGV PLEDFGHGHP
DPNLTYAKDL VDVMPDFGAA SDGDGDRNMV LGNFFVTPSD SVAIIAANAA IPYFRAGPKG
LARSMPTSGA LDRVAEKLKL PFFEVPTGWK FFGNLMDAGK LSICGEESFG TGSDHIREKD
GIWAVLAWLS ILAHRNVADV VKEYWATYGR NFFSRYDYEE CESEGANKMI EYLREILSNY
VLQFADDFSY TDPVDGSVAS KQGVRFVFTD GSRIIFRLSG TGSAGATVRI YIEQFEPDVS
KHDVDAQIAL KPLIDLALSV SKLKDFTGRE KPTVIT
>HsPGM1
TQAYQDQKPG TSGLRKRVKV FNYAENFIQS IISTVEPAQR QEATLVVGGD GRFYMKEAIQ
LIARIAAANG IGRLVIGQNG ILSTPAVSCI IRKIKAIGGI ILTASHNPGG PNGDFGIKFN
ISNGGPAPEA ITDKIFQISK TIEEYAVCLK VDLGVLGKQQ FKPFTVEIVD SVEAYATMLR
SIFDFSALKE LLSGPLKIRI DAMHGVVGPY VKKILCEELG APANSAVNCV PLEDFGGHHP
DPNLTYAADL VETMHDFGAA FDGDGDRNMI LGKFFVNPSD SVAVIAANII PYFQQTGVRG
FARSMPTSGA LDRVASATKI ALYETPTGWK FFGNLMDASK LSLCGEESFG TGSDHIREKD
GLWAVLAWLS ILATRKVEDI LKDHWQKYGR NFFTRYDYEE VEAEGANKMM KDLEALMFVY
TVEKADNFEY SDPVDGSISR NQGLRLIFTD GSRIVFRLSG TGSAGATIRL YIDSYEKDVA
KINQDPQVML APLISIALKV SQLQERTGRT APTVIT

### Material S1 The sequences used in phylogenetic analysis of PGM

>MpAPS1
VLGIILGGGA GTRLYPLTKK RAKPAVPLGA NYRLIDIPVS NCINSNVQKI YVLTQFNSAS
LNRHLSRAYG FVEVLAAQQS PEPNWFQGTA DAVRQYLWLF EENVLEFLVL AGDHLYRMDY
QNFIQAHRDT NADITVAALP EAFGLMKINE KGRIIEFAEK PSMGIYVVSK DAMIKLLRDD
FPANDFGSEV IPYWEDIGTI EAFYNANLGL TKFSFYDRTS PIYTQARFLP PSKMLDADDS
VIGEGCVIKN CKIYHSVVGL RSWIAEGAIV EDALLMGPMG IGRNSIIKRA IIDKNARIGE
NVKIGYFIKS GIVTIIKDAV IPNDTVI
>AtAPS2
VAAIVFGGGS DSELYPLTKT RSKGAIPIAA NYRLIDAVIS NCINSGITKI YAITQFNSTS
LNSHLSKAYR FVEVIAAYQS LEQGWFQGTA DAIRRCLWVF EEPVTEFLVL PGHHLYKMDY
KMLIEDHRRS RADITIVGLS FGFGFMEVDS TNAVTRFTIK GSAGIYVIGR EQMVKLLREC
LISKDLASEI IPYWEDVRSI GAYYRANMES IKYRFYDRQC PLYTMPRCLP PSSMSVAVNS
IIGDGCILDK CVIRGSVVGM RTRIADEVIV EDSIIVGRIG IGEKSRIRRA IVDKNARIGK
NVMIGYVIRE GIIIILRNAV IPNDSIL
>AtAPL3
VAAIILGGGD GAKLFPLTKR AATPAVPVGG CYRMIDIPMS NCINSCINKI FVLTQFNSAS
LNRHLARTYG FVEVLAATQT PGKKWFQGTA DAVRKFLWVF EDNIENIIIL SGDHLYRMNY
MDFVQHHVDS KADITLSCAP SEYGLVNIDR SGRVVHFSEK PSMGVYCFKT EALLKLLTWR
YPSNDFGSEI IPYWEDIGTI KSFYEANIAL VEFEFYDQNT PFYTSPRFLP PTKTEKCRNS
VISHGCFLGE CSIQRSIIGE RSRLDYGVEL QDTLMLGPIG IGRDTKIRKC IIDKNAKIGK
NVVIGFYIRS GITVVVEKAT IKDGTVI
>AtAPL2
VASIILGGGA GTRLFPLTSK RAKPAVPIGG CYRLIDIPMS NCINSGIRKI FILTQFNSFS
LNRHLSRTYG FVEVLAATQT SGKKWFQGTA DAVRQFIWVF EDNVEHVLIL SGDHLYRMDY
MNFVQKHIES NADITVSCLP SDFGLLKIDQ SGKIIQFSEK PSMGVYVFRK EVLLKLLRSS
YPSNDFGSEI IPYWEDIGTI GSFFDANLAL TEFQFYDQKT PFFTSPRFLP PTKVDKCRDS
IVSHGCFLRE CSVQHSIVGI RSRLESGVEL QDTMMMGPVG VGQNTKIKNC IIDKNAKIGK
NVVIGFHIRS GITVVLKNAT IRDGLHI
>MpAPL3
VACMILGGEA GSRLFPLTKR RAKSAVPMGG AYRLIDIQMS NCINSGINKV YVLTQFNSAS
LNRHISRTYG FVEVLAATQT LGKRWFMGTA DAVRRFSWIF DNAVEHVLVL SGDHLYRMNY
MDLVQSHHNS GADITVSCVP SAGGLLRLDH KGRVLSIHDK ASMGLYVFKK DVLIKLLKWI
YPSNDFASEI IPYWQDVGSI ESYFEANLAL TEFEFYDVSN PIYTSPRYLP PTTVDHCRDS
IVSHGCFLRS CSVQHSIIGI RSRIETGVEL KDVVMIGPMG VGEYSKIKRC IIDKNARIGK
NVILGIYIRS GIIIVSENAL IKDGMVI
>AtAPL4
VAAIILGGGN GAKLFPLTMR AATPAVPVGG CYRLIDIPMS NCINSCINKI FVLTQFNSAS
LNRHLARTYG FVEVLAATQT PGKKWFQGTA DAVRKFLWVF EDNIENILIL SGDHLYRMNY
MDFVQSHVDS NADITLSCAP SNFGLVKIDR GGRVIHFSEK PSMGVYCFKT EALLNLLTRQ
YPSNDFGSEV IPYWEDIGTI KTFYEANLAL VEFEFYDPET PFYTSPRFLP PTKAEKCRDS
IISHGCFLRE CSVQRSIIGE RSRLDYGVEL QDTLMLGPIG IGKDTKIRKC IIDKNAKIGK
NVIIGFYIRS GITVIVEKAT IQDGTVI
>E_coli
SVALILAGGR GTRLKDLTNK RAKPAVHFGG KFRIIDFALS NCINSGIRRM GVITQYQSHT
LVQHIQRGWE FVDLLPAQQR MKENWYRGTA DAVTQNLDII RRKAEYVVIL AGDHIYKQDY
SRMLIDHVEK GARCTVACMP SAFGVMAVDE NDKIIEFVEK PSMGIYVFDA DYLYELLEED
DRSHDFGKDL IPYWRDVGTL EAYWKANLDL ASLDMYDRNW PIRTYNESLP PAKFVQDRNS
LVSGGCVISG SVVVQSVLFS RVRVNSFCNI DSAVLLPEVW VGRSCRLRRC VIDRACVIPE
GMVIFYRSEE GIVLVTREML RKLGHKQ
>MpAPL2
VASLILGGGA GTRLFPLTRR RAKPAVPIGG AYRLIDVPMS NCINSGINKV FILTQFNSAS
LNRHLARTYG FVEVLAATQT PGMNWFMGTA DAVRQFTWLF EDGVEHVLIL SGDHLYRMDY
MDFVQKHKDS GADITISCVP SDYGLMKIDH KGQVLYFNEK PSMGIYVFKK EILLKLLRWR
YPANDFGSEI IPYWEDIGTI KSFFDANLAL TEFKFYDVAK PIFTSPRYLP PTKVEKCRDS
IVSHGCFLRD CSVEHSIVGI RSRIESGVEL QDTMMMGPLG VGTNSKIRNC IIDKNSRIGR
NVIIGFYIRS GITVVLKNST IKDGMVI
>AtAPS1
VLGIILGGGA GTRLYPLTKK RAKPAVPLGA NYRLIDIPVS NCLNSNISKI YVLTQFNSAS
LNRHLSRAYG FVEVLAAQQS PEPNWFQGTA DAVRQYLWLF EENVLEYLIL AGDHLYRMDY
EKFIQAHRET DADITVAALP TAFGLMKIDE EGRIIEFAEK PSMGIYVVSR DVMLDLLRNQ
FPANDFGSEV IPYWEDIGTI EAFYNANLGI TKFSFYDRSA PIYTQPRYLP PSKMLDADDS
VIGEGCVIKN CKIHHSVVGL RSCISEGAII EDSLLMGPIG IGKNSHIKRA IIDKNARIGD
NVKIGYFIKS GIVTVIKDAL IPTGTVI
>AtAPL1
VASIILGGGA GTRLFPLTKR RAKPAVPIGG AYRLIDVPMS NCINSGINKV YILTQYNSAS
LNRHLARAYG YVEVLAATQT PGKRWFQGTA DAVRQFHWLF EDDIEDVLIL SGDHLYRMDY
MDFIQDHRQS GADISISCIP SDFGLMKIDD KGRVISFSEK PSMGVYVFKK EILLNLLRWR
FPANDFGSEI IPYWEDIGTI RSFFEANLAL TEFSFYDAAK PIYTSRRNLP PSKIDNSKDS
IISHGSFLTN CLIEHSIVGI RSRVGSNVQL KDTVMLGPIG IGENTKIQEC IIDKNARVGK
NVIIGFYIRS GITVILKNSV IKDGVVI
>MpAPL1
VVSLILGGGA GTRLFPLTKR RAKPAVPIGG GYRLIDVPMS NCINSGINKV FILTQFNSAS
LNRHLARTYG FVEVLAATQT PGKEWFQGTA DAVRQYLWLF EDNLEDVLIL SGDHLYRMDY
MDFVEKHRNS GADITISCVP SDYGLMKIDD TGRVLYFSEK PSMGIYVFKK EILQKLLRWR
YPANDFGSEI IPYWEDIGTI KSFFDANLGL TAFSFYDAVK PIFTSPRYLP PSKIEKCRDS
IISHGCFLRD CSIKHSIVGI RSQMASGSAL KDTMMLGPIG VGANCKISNC IIDKNARIGS
NVVIGYYIRS GIVVILKNST IAPGTVI

Material S2 The sequences used in phylogenetic analysis of APL
